# The dual trxG/PcG protein ULTRAPETALA1 modulates H3K27me3 and directly enhances POLYCOMB REPRESSIVE COMPLEX 2 activity for fine-tuned reproductive transitions

**DOI:** 10.1101/2024.10.21.619451

**Authors:** Vangeli Geshkovski, Julia Engelhorn, Jean-Baptiste Izquierdo, Hamida Laroussi, Caroline Thouly, Laura Turchi, Meredith Wouters, Marie Le Masson, Emmanuel Thévenon, Ambre Petitalot, Lauriane Simon, Manon Verdier, Sophie Desset, Philipp Michl-Holzinger, Astrid Bruckmann, Hugues Parrinello, Klaus D. Grasser, Aline V. Probst, Raphaël Margueron, Gilles Vachon, Jan Kadlec, Cristel C. Carles

**Affiliations:** Grenoble Alpes University, CNRS, INRA, CEA, Plant and Cell Physiology Lab (LPCV), Bioscience and Biotechnology Institute of Grenoble, CEA, 17 rue des Martyrs, bât. C2, 38054 GRENOBLE Cedex 9, France; DIADE, Université de Montpellier, IRD, CIRAD, Montpellier, France; Grenoble Alpes University, CNRS, CEA, IBS, F-38000 Grenoble, France; Institut Curie, Paris Sciences et Lettres Research University INSERM U934/CNRS UMR3215, Paris, France; Institute of Genetics Reproduction and Development (iGReD), Université Clermont Auvergne, CNRS, Inserm, Clermont-Ferrand, F-63000, France; Cell Biology and Plant Biochemistry, University of Regensburg, D-93053 Regensburg, Germany; Biochemistry I, Regensburg Center for Biochemistry, University of Regensburg, D-93053 Regensburg, Germany; Cell MGX-Montpellier GenomiX, Univ. Montpellier, CNRS, INSERM, Montpellier, France

## Abstract

**Abstract:** The antagonistic POLYCOMB (PcG) REPRESSIVE COMPLEX 2 (PRC2) and trithorax (trxG) chromatin machineries play a major role in orchestrating gene expression during the development of multicellular eukaryotes. These complexes are well known for depositing and maintaining the repressive H3K27me3 and activating H3K4me3 marks, respectively. However, the mechanisms that govern the switch between these functions remains elusive, especially in plants, whose lifelong, flexible development relies heavily on this process. Here we demonstrate that the plant specific ULTRAPETALA1 (ULT1) protein, previously reported as a trxG factor that antagonizes the PRC2 enzymatic subunit CURLY LEAF (CLF), also exhibits a repressive function, increasing H3K27me3 levels at over a thousand genes. We discovered a physical interaction between ULT1 and PRC2 components, particularly the SWINGER (SWN) enzymatic subunit. We further show that *in vitro* ULT1 significantly enhances the enzymatic activity of PRC2^SWN^, and to a lesser extent also that of PRC2^CLF^, corroborating our epigenomic and developmental genetic data that reveal different ULT1 activity depending on the catalytic subunit of the PRC2 complex. This study provides new insights into the relative activities of CLF and SWN and introduces a novel mechanistic framework for a chromatin switch mediated by a bivalent trxG/PcG factor.

**Key message:** ULTRAPETALA1 counteracts or promotes PRC2 activity at hundreds of developmental genes in *Arabidopsis thaliana*, and activates the deposition of the repressive H3K27me3 chromatin mark via direct interaction with PRC2.

This is the first instance of a bivalent factor which functions as a cofactor of PRC2 HMTs.

---

Chromatin structure is intimately connected with gene expression and cell identity (Perino and Veenstra, 2016). Establishment of cell fates during development is accompanied by large-scale chromatin remodelling, which contributes to both lineage specification and maintenance. These epigenetic processes establish a chromatin framework that regulates the various developmental phases of multicellular eukaryotes.

Two antagonistic post-translational modifications carried by lysine residues of histone H3, H3K27me3 and H3K4me3, are key regulatory chromatin marks for genes of all higher eukaryotes (Kingston & Tamkun, 2014a; Mozgova & Hennig, 2015; Aranda *et al*., 2015); Schuettengruber *et al*., 2007). The Polycomb group (PcG) REPRESSIVE COMPLEX 2 (PRC2) and trithorax group (trxG) are responsible for the deposition of repressive and active trimethyl marks on H3K27 and H3K4, respectively. H3K27me3, in particular, is found at thousands of developmental genes, many of which are involved in cell fate determination, including differentiation and maintenance of cell identity. In animals, loss of PRC2 function leads to aberrant gene expression often affecting the expression of *HOX* genes and impairing stem cells differentiation, as initially reported in Drosophila (Schuettengruber *et al*., 2017). In plants, PRC2-mediated repression of MADS genes is essential for body and organ patterning, as well as developmental phase changes, throughout the lifespan. This is illustrated by the loss-of-function Arabidopsis mutants which display homeotic phenotypes and shifts in growth transitions (Engelhorn *et al*., 2014). Until now the antagonism between PcG/PRC2 and trxG functions has been demonstrated unidirectionally through the suppression of PRC2 mutant phenotypes by mutations in trxG genes (Berr *et al*., 2011; Kingston & Tamkun, 2014b).

Two PRC2 complexes are known in animals, containing the Enhancer of Zeste (E(z)) homolog 2 (EZH2) or the EZH1 enzyme, respectively, which catalyse di- or trimethylation of H3K27 (Margueron & Reinberg, 2011). Interestingly, EZH1 exhibits a lower enzymatic capacity than EZH2 (Margueron *et al*., 2008) (Lee *et al*., 2018). In Arabidopsis, there are three E(z) homologous PRC2 enzymatic subunits: one predominantly gametophytic, MEDEA (MEA), and two sporophytic, CURLY LEAF (CLF) and SWINGER (SWN), which catalyse all three levels of H3K27 methylation (Jacob *et al*., 2014; Borg *et al*., 2020). *In vitro* methylation assays have shown that MEA and CLF are active, however no data is available for SWN (Schmitges *et al*., 2011; Jacob *et al*., 2014; Borg *et al*., 2020).

The apparent functions of the PRC2 sporophytic enzymes CLF and SWN, as revealed by loss-of-function mutants, are not equivalent: *clf* mutants show upward leaf curling, early flowering and flowers with homeotic defects. These phenotypes are largely caused by ectopic expression of the MADS floral genes *AGAMOUS (AG)* and *APETALA3* (*AP3)* (Goodrich *et al*., 1997). Interestingly, although the *FLOWERING LOCUS C* (*FLC*) MADS gene, central repressor of flowering, is highly upregulated in *clf* mutants, these plants flower early, showcasing the complexity of the pleiotropic phenotypes of *prc2* mutants. For *swn* mutants, plants are not clearly distinguishable from wild-type plants except for a slight flowering time defect (Shu *et al*., 2019). This was also observed at the molecular level, as very few genes show a change in H3K27me3 or transcript levels in *swn* compared to wild-type plants (Shu *et al*., 2019). *clf swn* double mutants, however, develop callus-like masses of disorganized tissue after normal embryonic development. These structures contain a multitude of tissue types resulting from uncontrolled cell differentiation and due to ectopic de-repression of thousands of developmental genes (Chanvivattana *et al*., 2004). Of note, the differential impacts of CLF and SWN on development and gene regulation cannot be explained by different occupancies at the chromatin as their genome-wide binding profiles largely overlap (Shu *et al*., 2019). This raises questions on the differential roles of CLF and SWN in depositing the H3K27me3 mark at the same loci and on their functional interplay (Shu *et al*., 2020).

The PRC2 complex of Drosophila comprises three other sub-units: Extra Sex Comb (ESC) or its homolog ESCL; Suppressor of Zeste 12 (SU(Z)12) and p55 (Czermin *et al*., 2002). While there is only one ESC homolog in Arabidopsis, FERTILIZATION INDEPENDENT ENDOSPERM (FIE), there are three SU(Z)12 homologs: EMBRYONIC FLOWER 2 (EMF2), VERNALIZATION INSENSITIVE 2 (VRN2) and FERTILIZATION-INDEPENDENT SEED 2 (FIS2); and five p55 homologues: MULTICOPY SUPPRESSOR OF IRA 1-5 (MSI1-5, with only MSI1 demonstrated to be part of the PRC2 complex).

The regulation of PRC2 enzymatic activity in mammals has attracted considerable attention. It is now well established that a set of cofactors interact with SU(Z)12 and are essential for the proper targeting and activity of PRC2 (Højfeldt *et al*., 2019). While most of these cofactors have paralogs in Drosophila, the situation appears quite different in plants, where many PRC2 direct interactors are unique to the green lineage (Godwin & Farrona, 2022). Moreover, the impact and mechanism of these plant accessory proteins on the activities of PRC2 remain largely unknown.

We previously identified a PRC2 antagonist, called ULTRAPETALA1 (ULT1), that promotes developmental switches such as floral transition and flower morphogenesis in *Arabidopsis thaliana*. We found that ULT1 regulates stem cell maintenance and differentiation in the shoot apex, via transcriptional regulation of developmental loci (Carles *et al*., 2004, 2005). When ectopically expressed in plants, it induces homeotic phenotypes through the ectopic expression of meristem and flower MADS genes and reduction in H3K27me3 levels at these loci. Moreover, when combined with a mutation in the PRC2 catalytic subunit CLF, *ult1* loss-of-function hides PRC2 mutant phenotypes (Carles & Fletcher, 2009). This trxG-type of function of ULT1 was explained by the direct regulation of *AG* and *AP3* (Carles & Fletcher, 2009).

On the other hand, ULT1 was also reported to have a Polycomb-type function. Indeed, ULT1 together with the trxG ATX1 and PcG EMF1 members, prevent seed gene expression after germination (Xu *et al*., 2018). This contribution to seed gene repression in seedlings correlates with a decrease in H3K27me3 levels, only observed in the specific genetic context of the *ult1 atx1 emf1* triple mutant. Thus, while ULT1 derepresses flower genes, reducing their H3K27me3 levels, it contributes to repression of seed genes, which associates with elevated H3K27me3 levels. These two sets of seemingly contrasting findings position ULT1 as a potential dual regulator of chromatin dynamics.

To further investigate ULT1 opposing regulatory activities on PRC2, we employed a genome-wide approach assessing the effect of ULT1 on the abundance of the H3K27me3 mark, using loss- and gain-of-function lines. We found that, on one hand, ULT1 reduces H3K27me3 amounts at over a fourth of CLF targets, and that these are enriched in transcriptional and developmental functions. Interestingly, the genes de-repressed by ULT1 poorly overlap with targets of the H3K27me3 Histone demethylases (HDMT) REF6, ELF6 and JMJ13. On the other hand, ULT1 promotes an increase in H3K27me3 amounts at a third of genes known to be repressed by the other sporophytic H3K27 histone methyltransferase (HMT), SWN. We found that ULT1 physically interacts with both CLF and SWN, with a better affinity for SWN, and does not interfere with the PRC2 complex assembly. Moreover, we also show that ULT1 significantly increases the *in vitro* HMT activity of the SWN-containing PRC2 complex and to a lesser extent that of the CLF-containing PRC2 complex. Finally, phenotypic and molecular analyses of flowering time and flower organ number lead to a consistent interaction with PRC2, along with the co-regulation of key developmental genes.

## Results

### ULT1 has a dual genome-wide effect on H3K27me3 abundance

To assess the effect of ULT1 on PRC2-induced H3K27me3 deposition, we quantified global and genome-wide levels of H3K27me3 in the *ult1-2* and *ult1-3* loss-of-function lines (Carles *et al*., 2005), thereafter referred to as *ULT1* lof, as well as in a *35S::ULT1* gain-of-function line (Carles & Fletcher, 2009), thereafter referred to as *ULT1* gof. These experiments revealed that ULT1 has a dual effect on H3K27me3 abundance, inducing both its decrease and increase at two distinct sets of genes.

#### ULT1 negatively regulates H3K27me3 enrichment at 916 loci

Chromatin immuno-precipitation assays followed by sequencing (ChIP-seq) revealed that 2925 genes show a consistent decrease of sequencing-depth-normalized H3K27me3 abundance in *ULT1* gof (genes overlapping with regions of significantly reduced H3K27me3 levels in both replicates, for details see methods and Suppl. Fig.S1), while 1532 and 1378 show an increase in *ult1-2* and *ult1-3* lof (genes overlapping with regions of significantly increased H3K27me3 levels in both replicates), respectively (Fig. 1A). 916 genes are shared between these three sets (Table S1), which we named ULT1_derep, as they are target genes for de-repression by ULT1. The ULT1_derep population shows an enrichment in the Gene Ontology (GO) development-related processes and transcription factors (Fig. 1B and Suppl. Fig.S1), including meristem, flowering and flower morphogenesis regulators. Among these are the flower MADS transcription factor-encoding genes, in particular *AGAMOUS* (*AG*), *APETALA3* (*AP3*) and *SEPALLATA1-4* (*SEP1-4*), previously reported to be targets for activation by ULT1 (Carles & Fletcher, 2009). In addition to these already known targets, we found *AP1*, *CAULIFLOWER* (*CAL*), *PISTILLATA* (*PI*) and the flowering regulators *AGAMOUS-LIKE 24* (*AGL24*) and *FLOWERING LOCUS T* (*FT*) (Fig. 1C; Suppl. Fig. S2; Table S1). This dataset thus supports a model where ULT1 functions as a PRC2 antagonist.

**Figure 1.**
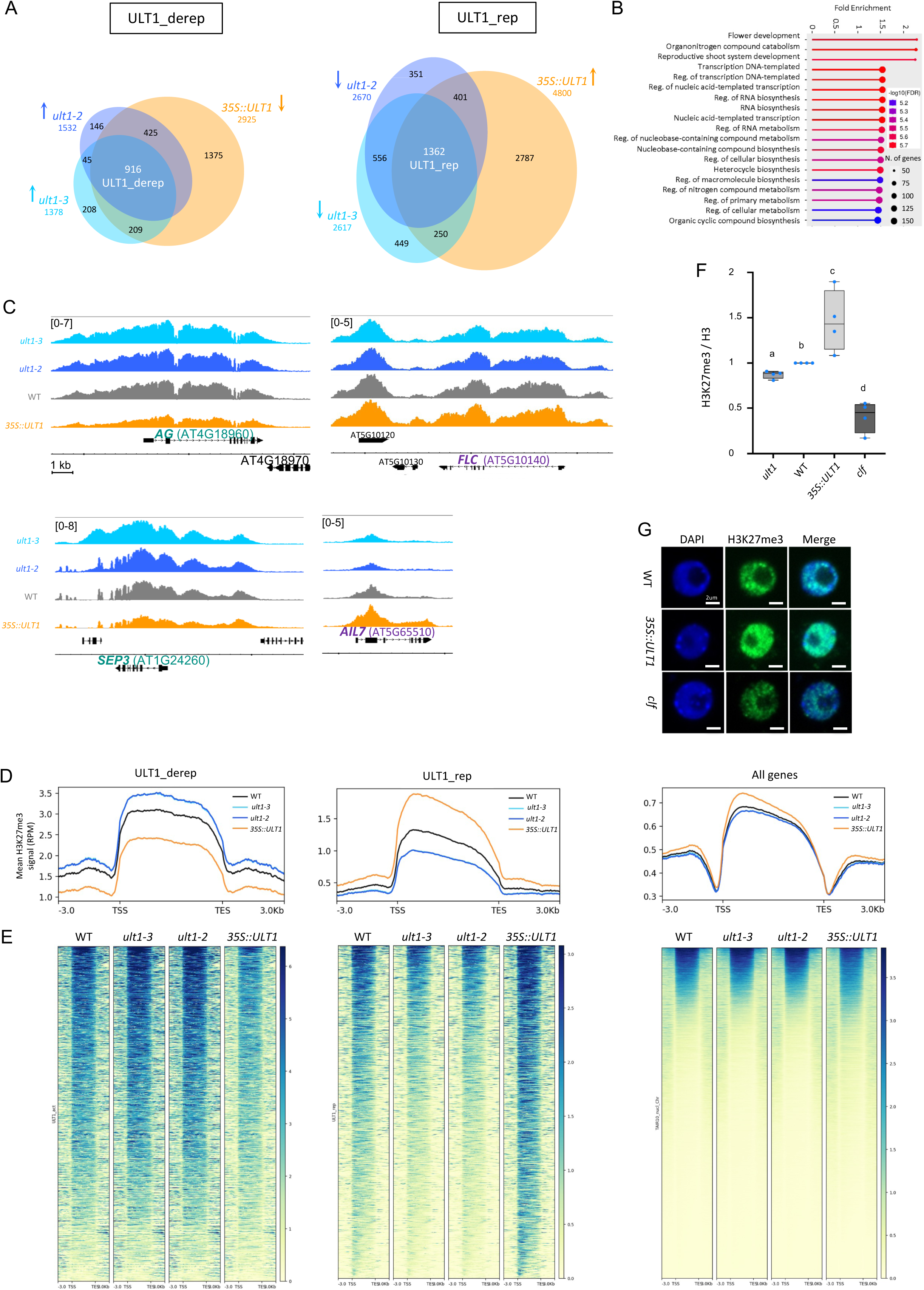
Global and genome-wide dual effect of ULT1 on H3K27me3 mark abundance. **(A-E)** Genome-wide influence of ULT1 on H3K27me3 marks in 8 day-old seedlings (grown in long-day conditions at 18°C day / 16°C night) of ULT1 loss- and gain-of-function lines: WT (Ler wild type), *ult1-2* (EMS line, null mutant), *ult1-3* (T-DNA insertion line, null mutant) and *35S::ULT1* (ectopic over-expressor). **(F,G)** Global effect of ULT1 gain-of-function on H3K27me3 abundance in 8 day-old seedlings, assessed by western blot or *in cyto* immuno-fluorescence. **(A)** Venn-diagram showing overlap between populations of genes (left) with significantly elevated H3K27me3 in *ult1* mutants and significantly reduced H3K27me3 in *35S::ULT1* (ULT_derep); and (right) with significantly reduced H3K27me3 in *ult1* mutants and significantly increased H3K27me3 in 35S::ULT1 (ULT1_rep). Numbers in the circles indicate the number of genes in the respective overlap, numbers under genotypes indicate the total number of significantly changed genes found in this genotype. Significance of overlap between all set of genes were calculated one-to-one using a Fisher’s test, and all were found to be lower than 2.2e-16. **(B)** Functional categorisation of ULT1_derep genes. ShinyGO v0.8 analysis for significantly over-represented Gene Ontology terms. **(C)** Examples of influence of ULT1 on H3K27me3 marks. Genome browser (IGB) view of H3K27me3 determined by ChIP-seq in wild type, *ult1-2*, *ult1-3* and *35S::ULT1* lines at the (left) *AGAMOUS* (*AG*) and *SEPALLATA3* (*SEP3*) loci (examples of ULT_derep targets) and at the (right) *FLOWERING LOCUS C* (*FLC*) and *AINTEGUMENTA-LIKE 7* (*AIL7*) loci (examples of ULT_rep targets). Thick arrows indicate exons, thin lines represent introns. The min-max coverage values for each gene are given in brackets (same scale for all genotypes corresponding to each gene). Note that the oscillation of the data towards the 3’-end of the *SEP3* locus is caused by the mapping of (Ler) ecotype to the Columbia reference genome. *ult1-3* does not show this alteration since it was originally generated in Columbia and backcrossed. **(D)** Metagenes for ULT1-derep, ULT1-rep genes, and all genes: profile of H3K27me3 in WT, *ult1-2*, *ult1-3* and *35S::ULT1* lines. Mean distribution of H3K27me3 over ULT1-derep, ULT1-rep genes, and genome-wide. Genes are scaled to the same length (6kb), Panels represent data of one biological replicate, similar observations were made for the second replicate. RPM: reads per million. **(E)** Heatmaps for H3K27me3 distribution for all ULT1-derep, ULT1-rep genes, and genome-wide, ranked from (top line) the highest H3K27me3-marked to(bottom line) the lowest H3K27me3-marked gene in the WT. **(F)** Quantification of H3K27me3 by western blot on 8 day-old seedlings (*ult1-3*, WT, *35S::ULT1* and *clf-2*), represented as box plots. Mean was calculated from 4 replicates (which values are indicated by dots), for which H3K27me3 quantification was normalized to H3. Letters indicate significant differences between samples (Mann-Whitney test, P-value < 0.05). **(G)** Representative nuclei after immunofluorescence staining using an antibody directed against H3K27me3 in WT, *35S::ULT1* and *clf-2* seedlings. DNA was counterstained with DAPI.

#### ULT1 positively regulates H3K27me3 enrichment at 1362 loci

Remarkably, the ChIP-seq data also revealed an opposite effect of ULT1 on H3K27me3 abundance, demonstrating that ULT1 functions also as a potent PRC2 agonist. Indeed, in the same analysis as described above, 4800 genes showed an increase in H3K27me3 in *ULT1* gof, and 2670 and 2617 showed a decrease in *ult1-2* and *ult1-3* lof, respectively (Fig. 1A). The 1362 genes common to these three sets were named ULT1_rep (genes repressed by ULT1) (Table S1). GO analyses do not reveal enrichment in development-related processes and transcription factors terms, even though the ULT1_rep genes comprise some flowering time and meristem regulators. These include several developmental genes coding for transcription factors, including the MADS gene *FLOWERING LOCUS C* (*FLC*), master regulator of the reproductive transition and its two paralogues *MADS AFFECTING FLOWERING 4* (*MAF4*) and *MAF5*, *AGAMOUS-LIKE LOCUS 51* (*AGL51*), the meristem regulators *AINTEGUMENTA-LIKE 7* (*AIL7*), *CUP-SHAPED COTYLEDON 1* (*CUC1*), *ARABIDOPSIS PIN-FORMED 7* (*PIN7*) and *PERIANTHIA* (*PAN*) (Fig. 1C; Suppl. Fig. S2; Table S1). In contrast to previously reported biologically relevant ULT1_derep genes (Carles & Fletcher, 2009; Pu *et al*., 2013), there is little evidence for ULT1 function as a repressor, except in the context of seed germination (Xu *et al*., 2018). We therefore confirmed the effect of ULT1 through ChIP-qPCR, on some of the ULT1_rep genes (Suppl. Fig. S3). Together, these results reveal that, in addition to reducing H3K27me3 levels at many genes, ULT1 can also enhance the accumulation of this mark across a broad set of genes.

#### ULT1 impacts H3K27me3 throughout target loci and leads to overall H3K27me3 increase in the nucleus

For both ULT1_derep and ULT1_rep genes, mark abundance changes can be observed throughout the target loci (Fig. 1C-E; Suppl. Fig. S2). Hence, ULT1 impacts the entire H3K27me3-marked region of genes it regulates.

Moreover, meta-gene analyses reveal an overall promoting function of H3K27me3 abundance by ULT1 (summed-up effect on all targets) (Fig. 1D, E). We assessed if the global changes in mark detected by ChIP-seq could be observed by immunodetection in 8-day-old seedlings. Quantifications from immunoblots reproducibly revealed changes in H3K27me3 levels, higher in *35S::ULT1* and lower in *ult1-3* plants, along with the expected loss of H3K27me3 in the *clf-2* mutant (Fig. 1F; Suppl. Fig. S4; Suppl. Fig. S5). These results were consistent with immunofluorescence staining, which was initially performed to compare the global H3K27me3 pattern within the nucleus across genotypes. While the nuclear H3K27me3 distribution was not detectably altered, *35S::ULT1* samples consistently and reproducibly exhibited stronger signals than WT, whereas *clf-2* samples displayed the opposite trend (Fig. 1G).

#### ULT1 impacts fewer targets for H3K4me3 abundance and gene expression, yet changes occur for common flowering and flower morphogenesis genes

Next, we wanted to assess whether the changes we observed in H3K27me3 were correlated with changes in the abundance of the antagonistic mark H3K4me3. We thus analysed H3K4me3 genome-wide in the same ULT1 lof and ULT1 gof lines, using the same chromatin samples as those immuno-precipitated for the H3K27me3 ChIP-seq experiments. We also assessed the correlation with gene expression changes through RNA-seq experiments (Suppl.Tables S2 and S3) on the same tissues. We found that, compared to its effect on H3K27me3 levels, ULT1 has a less striking effect on the abundance of the H3K4me3 mark and on gene expression (Suppl. Fig. S2B,D; Suppl. Fig. S6 and S7). Indeed, only 45 genes show increase in the H3K4me3 mark in *ULT1* gof, and 6 genes show decrease in the H3K4me3 mark in both *ULT1* lof lines. On the other hand, 117 genes show decrease in the H3K4me3 mark in *ULT1* gof line, and 24 genes show increase in the H3K4me3 mark in both *ULT1* lof lines (Suppl. Fig. S6A). GO analyses reveal that the population of genes which lose the H3K4me3 mark in ULT1 gof is enriched in development and cell fate specification related terms (Suppl. Fig. S8). Transcriptional analyses showed that 14 genes were down-regulated and 48 were up-regulated in both ULT1 lof lines, while the *35S::ULT1* gof line had 136 up-regulated and 81 down-regulated genes (Suppl. Fig. S7A, B).

Likely as a consequence of their smaller sizes, the populations of genes with significantly changed H3K4me3 abundance or of differentially expressed genes, have a limited overlap with the populations of genes with significantly changed H3K27me3 abundance (Suppl. Fig. S6 and S7). Even without a threshold for significant differences in expression, only a very limited number of ULT1_derep/ULT1_rep genes displayed expression changes related to their H3K27me3 changes in the lof and gof lines. Indeed, a majority of genes were expressed at very low levels if at all, consistent with H3K27me3-targeting (Suppl. Fig. S7 C). Yet, some specific genes that are biologically relevant regarding ULT1 developmental functions, were shared between the H3K27me3, H3K4me3 and expression changing lists. Among the ULT1_derep genes (identified previously based on H3K27me3 changes), 6 also showed elevated H3K4me3 in *ULT1* gof, including *SEP3,* and 10 showed elevated expression in *ULT1* gof, including *SEP3*, *FT* and *PI* (Table S3). On the other hand, *FLC,* which is part of ULT1_rep genes, is one of the very few genes that also shows H3K4me3 increase in *ult1-2* lof and decrease in *35S::ULT1* gof, and significantly and coherently changes in expression in our transcriptome dataset, for all ULT1 lof and gof lines (Suppl. Fig. S2B, D and Suppl. Tables S2 and S3).

Together, these findings show that ULT1 modulates H3K27me3 levels, either enhancing or reducing them across broad gene sets, thereby acting as both an antagonist and agonist of PRC2.

### ULT1 de-represses mainly CLF targets, independently of JmJ Histone Demethylases

We and others previously reported that ULT1 counteracts the repressive activity of CLF (Carles & Fletcher, 2009; Pu *et al*., 2013). In order to assess the extent of ULT1 antagonistic effect on CLF, we compared H3K27me3 ChIP-seq datasets for both factors (Shu *et al*., 2019) and found that CLF targets for repression (thereafter referred to as ‘CLF_rep’, i.e. genes showing decrease in H3K27me3 amount in *clf* lof) are greatly over-represented among ULT1_derep genes (70,7%, of ULT1_derep are also CLF_rep, p-value<0.001; while only 0,3% of ULT1_derep are SWN_rep, i.e. genes showing decrease in H3K27me3 amount in *swn* lof). In agreement with these results, ULT1_derep are greatly over-represented among CLF_rep (23,6%, of CLF_rep are also ULT1_rep, p-value<0.001; while only 1,6% of SWN_rep are ULT1_derep) (Fig. 2A, Suppl. Table S4). These data indicate that the antagonistic function of ULT1 toward PRC2 is more related to CLF than to SWN.

**Figure 2.**
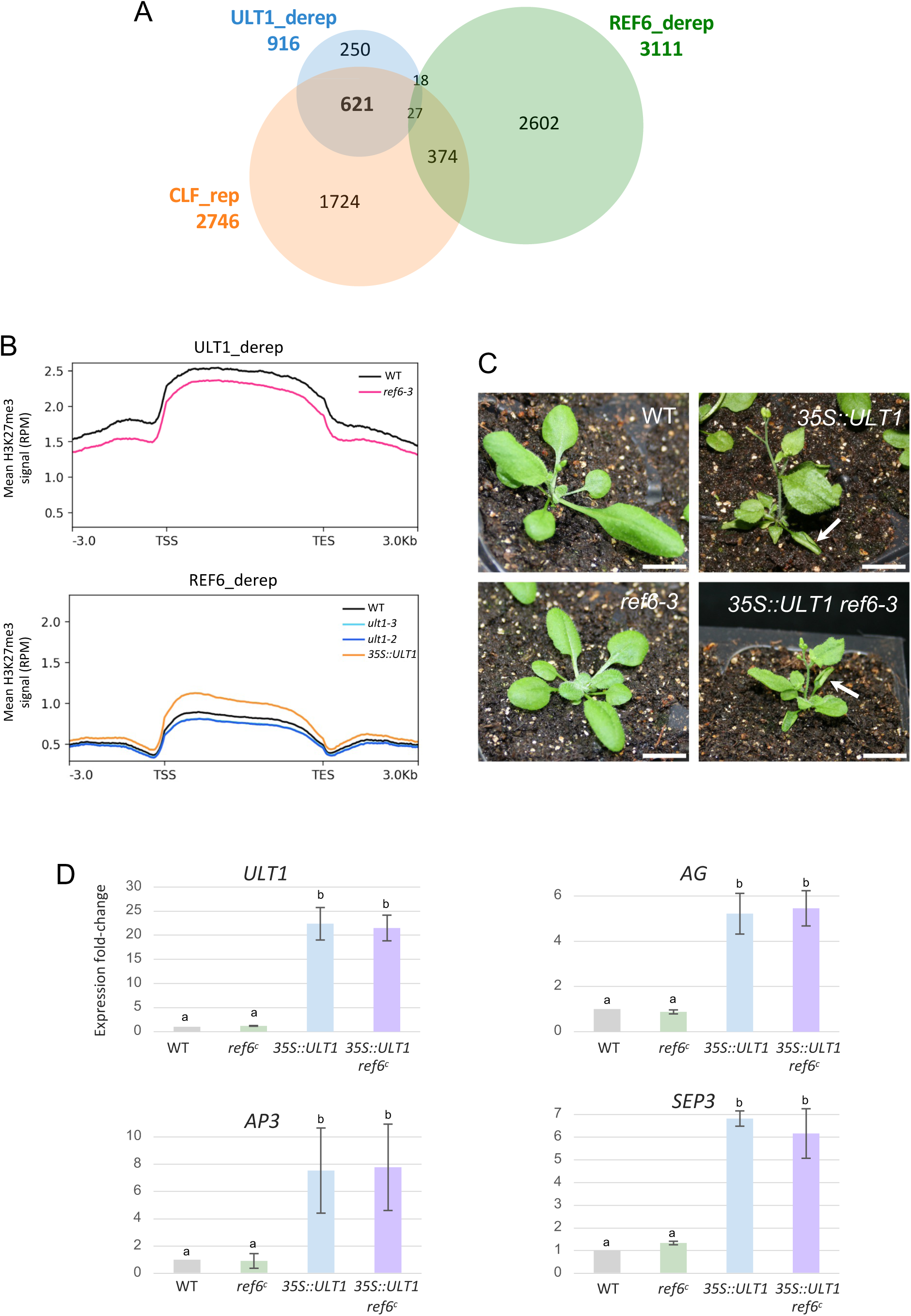
REF6 is dispensable for ectopic ULT1-induced phenotypes and de-repression of MADS floral genes. **(A)** Venn-diagram showing overlap between populations of ULT1_derep genes with CLF_rep genes (genes with significantly reduced H3K27me3 in *clf*, re-analysis of raw data from Shu *et al*. 2019) and REF6_act genes (genes with significantly elevated H3K27me3 in *ref6*). **(B)** Metagenes representing the mean H3K27me3 levels on genes de-repressed by ULT1, as observed in WT and *ref6* mutant (top), and the mean H3K27me3 levels on genes de-repressed by REF6, as observed in WT, *ult1-2, ult1-3* and *35S::ULT1* lines (bottom). RPM: reads per million. **(C)** Pictures of 28 day-old Col-0 plants (WT, *35S::ULT1*, *ref6-3* mutant, *35S::ULT1 ref6-3), i*ndicating that 35S::ULT1 ectopic expression phenotypes do not depend on REF6 function. Arrows point at upward curling leaves in *35S::ULT1* and *35S::ULT1 ref6-3* plants. **(D)** Relative expression levels of *ULT1*, along with the homeotic floral genes *AG, AP3* and *SEP3* in 8-day-old seedlings (F3 *ref6C* x *35S::ULT1*), assessed by RT-qPCR. Expression levels of target genes were normalized with the reference gene *MON1*. Error bars correspond to standard deviation for 3 technical replicates. Letters indicate significant differences between samples (Student’s t-test, one sided paired, P-value < 0.05). Similar results were obtained for 2 biological replicates.

Similarly to ULT1, the RELATIVE OF EARLY FLOWERING 6 (REF6) demethylase can partially counteract CLF activity (Lu *et al*., 2011) by reducing the amount of H3K27me3 on common genes. We thus wondered whether ULT1 could counteract CLF activity by decreasing H3K27me3 amount through the REF6 pathway (Yan *et al*., 2018), but found that ULT1_derep and REF6_derep populations share very few common genes. H3K27me3 levels are even changing in opposite directions when considering the overall abundance of the mark over all ULT_derep genes in *ref6* mutants and vice versa (Fig. 2A: 45 genes, corresponding to 1,45% of REF6_derep targets, p-value= 2,37.10^-121^ for under-representation, and corresponding to 4,91% of ULT1_derep genes, p-value= 0 for under-representation and Fig. 2B). This reveals that ULT1 largely acts in a REF6-independent manner to reduce H3K27me3 levels at target genes. Comparison of H3K27me3 ChIP-seq data between ULT1 and all JmJ HDMTs (analyses in triple mutants for REF6, ELF6 and JMJ12) (Yan *et al*., 2018), leads to similar conclusions (Suppl. Fig. S9-S11). Moreover, we constitutively expressed *ULT1* in the *ref6-3* mutant background (Fig. 2C) and found that *35S::ULT1* and *35S::ULT1 ref6-3* plants displayed similar developmental defects (small plants with upward curling leaves). We next analysed the expression levels of the MADS-box genes *AG*, *AP3* and *SEP3,* for which H3K27me3 levels are regulated by ULT1, CLF and REF6. We found that these three genes are ectopically expressed to similar levels in *35S::ULT1* and *35S::ULT1 ref6c* (Yan *et al*., 2018) seedlings (Fig. 2D), demonstrating that ULT1 induces de-repression independently from REF6, even at common target genes. Similarly, *35S::REF6-YFP-HA ult1-3* plants do not differ from *35S::REF6-YFP-HA* plants in appearance and both display ectopic expression of *AG*, *AP3* and *SEP3* (Suppl. Fig. S12). This shows that ULT1 is dispensable for ectopic REF6-dependent phenotypes and de-repression of MADS floral genes. Together, these results strongly indicate that ULT1 acts in a REF6-independent manner to decrease H3K27me3 amounts.

### ULT1 physically interacts with the core PRC2 subunits

To shed light onto the mechanism underlying the ULT1-mediated PRC2 regulation, we investigated potential interactions between ULT1 and PRC2 components in plant cells. To this end, we performed affinity purification coupled with mass spectrometry experiments, conducted on *Arabidopsis thaliana* cell cultures (Antosz *et al*., 2017) expressing a tagged ULT1 fusion protein (GS-ULT1). The results show that ULT1 copurifies in all three replicates with two core subunits of PRC2: SWN and FIE (Fig. 3A, Suppl. Table S5). In addition, we identified an association between ULT1 and ENHANCER OF LIKE HETEROCHROMATIN 1 (EOL1), which interacts physically with CLF and SWN to maintain the H3K27me3 mark in plants (Zhou *et al*., 2017). This data reveals that ULT1 can associate with the PRC2 complex in plant cells, potentially by direct interaction.

**Figure 3.**
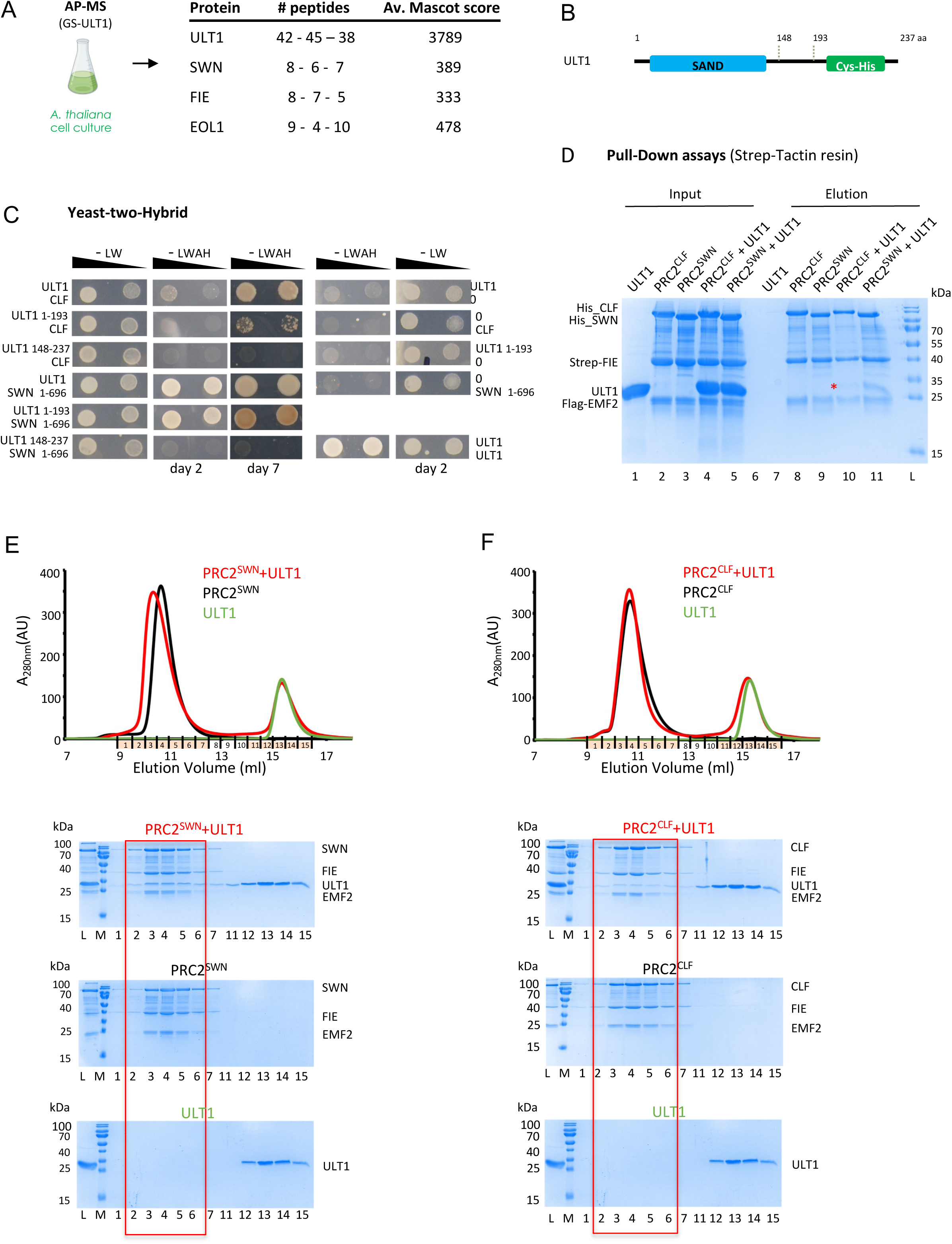
ULT1 physically interacts with PRC2 HMT CLF and SWN, *in vivo* and *in vitro*, with better affinity for SWN. **(A)** AP-MS using GS-ULT1 in *Arabidopsis thaliana* cell cultures reveals components of the PRC2 complex. The numbers of unique peptides obtained for each replicate and the average mascot score out of 3 replicates are given. **(B)** Schematic representation of ULT1 structure in domains. **(C)** Yeast-two-hybrid assays for interaction tests between ULT1 (full-length, ULT1_1-193_ and ULT1_148-237_) and CLF or SWN. Photographs were taken after 2 days or 7 days of yeast growth, at two concentrations corresponding to optical densities at 600 nm (OD) of 10 and 1. The ULT1-ULT1 interaction was used as positive control. **(D)** Strep-tag pull-down experiments of PRC2^SWN^ and PRC2^CLF^ complexes with untagged ULT1. Catalytic lobes of PRC2^SWN^ and PRC2^CLF^ contained a Strep-tag on the FIE subunit. All proteins were first purified by affinity chromatography and gel filtration. Proteins were mixed as indicated above the lanes. Input (lanes 1–5) and the eluates (lanes 7–11) were analyzed on 15% SDS-PAGE gels stained with Coomassie brilliant blue. The red asterisk indicates position of ULT1 in the elutions. PRC2^SWN^ copurifies more efficiently with ULT1 than PRC2^CLF^ (compare lanes 10 and 11). **(E)** Overlay of Superdex 200 gel filtration elution profiles of the catalytic lobe of PRC2^SWN^, ULT1 and their mixture. Lower panels show SDS-PAGE analysis of fractions 1-7 and 11-15 corresponding to the two major elution peaks. The red rectangle highlights fraction where ULT1 co-elutes with PRC2^SWN^. **(F)** Overlay of Superdex 200 gel filtration elution profiles of the catalytic lobe of PRC2^CLF^, ULT1 and their mixture. Lower panels show SDS-PAGE analysis of fractions 1-7 and 11-15 corresponding to the two major elution peaks. The red rectangle highlights fraction where ULT1 co-elutes with PRC2^CLF^. ULT1 is, however, sub-stoichiometric and there is essentially no shift of the elution peak compared to PRC2^CLF^alone. The elution profile and corresponding SDS-PAGE gel of ULT1 is the same in panel E and F.

To verify whether ULT1 interacts with components of PRC2 directly, we performed yeast-two-hybrid (Y2H) experiments and found that full-length (FL) ULT1 interacts with both FL CLF and SWN_1-696_, a C-terminally truncated version of SWN lacking its SET domain (Fig. 3B, C). Yeast growth on selective medium was however much faster with SWN (comparable to that observed for the already reported ULT1-ULT1 interaction (Moreau *et al*., 2016)). ULT1_1-193_, encompassing the N-terminal SAND domain was sufficient for the interactions with CLF and SWN, signifying that the C-terminal Cys-rich domain of ULT1 is not absolutely required. We detected no interaction between ULT1 and the core PRC2 protein FIE by Y2H (data not shown). These results strongly indicate that ULT1 can directly interact with both sporophytic HMTs of PRC2.

To test whether ULT1 can directly interact with the PRC2 complexes *in vitro*, we overexpressed in insect cells the catalytic lobes of both PRC2^CLF^ and PRC2^SWN^ containing either the full-length CLF or SWN, FIE and the VEFS domain of EMF2. We based these choices on the known atomic structures of the PRC2 complex in yeast and humans (Jiao & Liu, 2015; Justin *et al*., 2016). The purified PRC2 complexes were used in Strep-tag pull-down assays with pure full-length ULT1 (Fig. 3D, Suppl. Fig. S13). While PRC2^SWN^ bearing a Strep-tag on the FIE subunit could efficiently copurify ULT1 (Fig. 3D, lane 11), in the PRC2^CLF^ co-purification ULT1 was sub-stoichiometric (Fig. 3D, lane 10). The preferential interaction of ULT1 with PRC2^SWN^ was further confirmed by size exclusion chromatography, where ULT1 efficiently formed complex with PRC2^SWN^, while in complex with PRC2^CLF^, it co-eluted as sub-stoichiometric (Fig. 3E,F).

### ULT1 enhances PRC2 complex activity mainly through SWN, both *in vitro* and *in vivo*

Given the direct interaction of ULT1 with PRC2, we wondered whether this might impact the enzymatic activity of the complex. We first addressed this question with *in vitro* Histone Methyl Transferase (HMT) assays, monitoring recombinant PRC2 activity in the presence or absence of ULT1, using substrate nucleosomes that were either reconstituted from recombinant histones (bacterially expressed therefore seen as lacking post-translational modifications, Fig. 4A) or “native” histones purified from HeLa cells (carrying various post-translational modifications, Fig. 4B). ULT1 by itself has no enzymatic activity (Fig. 4A,B, lanes 1 to 3). PRC2^CLF^ displays a robust activity which appears rather insensitive to the presence of ULT1 on recombinant histone H3 and limited on H3 from cell extracts (Fig. 4A, B, lanes 4 to 7). In contrast, PRC2^SWN^, in these assays, has a modest basal activity which is strongly enhanced in the presence of ULT1 on recombinant substrate (up to 8-fold when the MBP tag of ULT1 is removed (Fig. 4A, B, lanes 8 to 11). On native substrate, PRC2^SWN^ and PRC2^CLF^ are less distinct, displaying a more similar level of basal activity, and while the enzymatic activities of both complexes are enhanced by ULT1, the effect appears slightly more pronounced on PRC2^SWN^. Importantly, while we used a molar excess of ULT1 as compared to PRC2 for these assays, we also performed a titration of ULT1 that showed that its stimulatory effect on PRC2 enzymatic activity can be observed with a much lower amount of protein (Suppl. Fig. S14). ULT1 thus seems to function as a strong co-activator of PRC2 with a clear preference for SWN in this assay. Importantly, close examination of the gel also reveals that CLF and SWN are automethylated as it has been described for mammalian PRC2(Lee *et al*., 2019; Wang *et al*., 2019) but ULT1 does not appear to be methylated by PRC2. Besides, ULT1 is unable to regulate the mammalian PRC2, which may reflect a plant specific mechanism of action (Suppl. Fig. S15).

**Figure 4.**
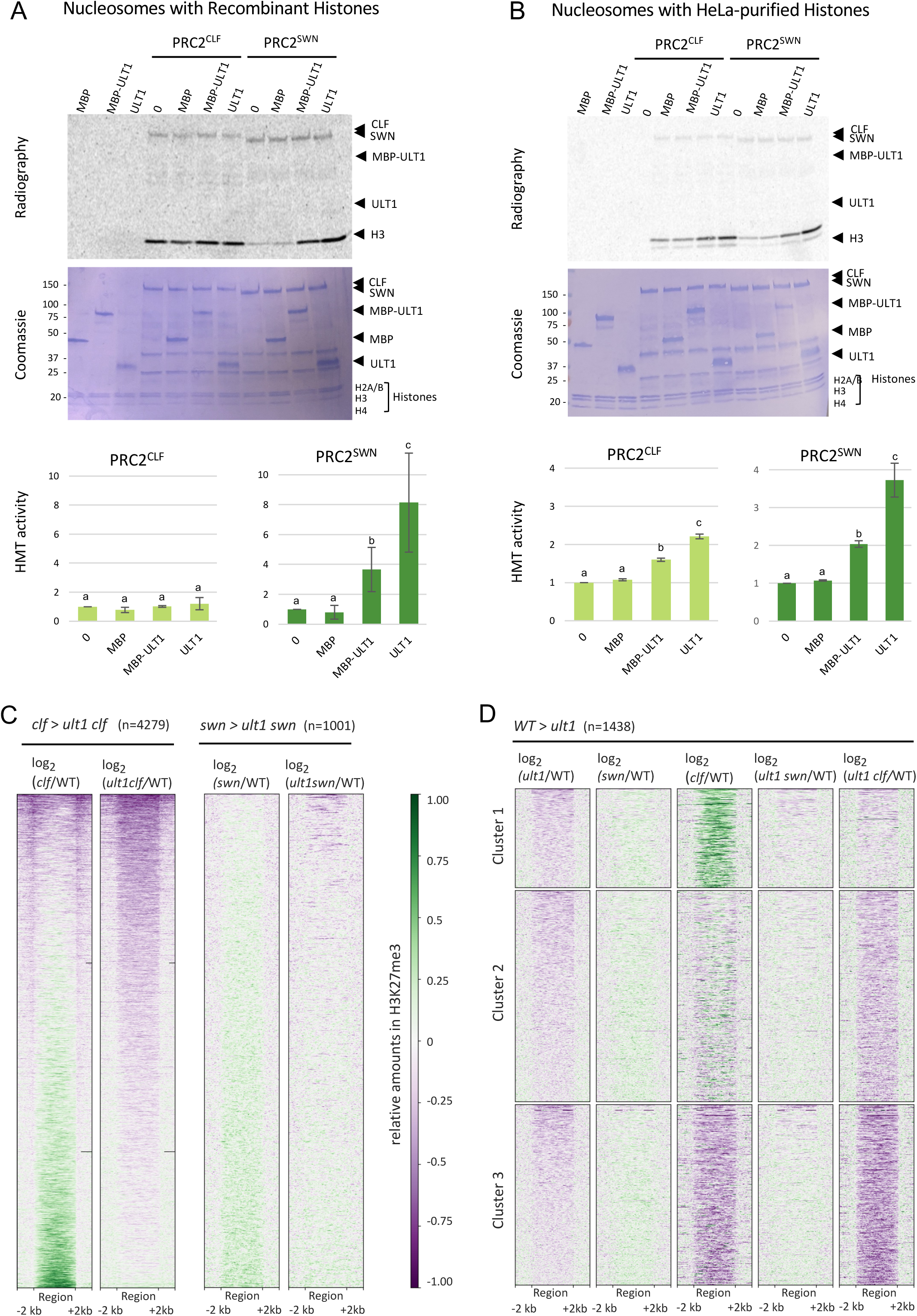
ULT1 increases PRC2 activity with stronger enhancement of PRC2^SWN^ function *in vitro* and *in vivo*. **(A-B)** ULT1 increases PRC2 HMT activity. *in vitro* HMT activity tests on nucleosomes prepared from **(A)** recombinant or **(B)** HeLa-purifed histones indicate that ULT1 significantly increases PRC2^SWN^ activity on non-modified histones and both PRC2^SWN^ and PRC2^CLF^ activities on HeLa-purified histones. Top panels show autoradiography of a SDS-PAGE gel after incubation of PRC2^CLF^ or PRC2^SWN^ (1.5ug of complex) without or with MBP, MBP-ULT1 or ULT1. Middle panels show Coomassie staining of the same SDS-PAGE. Bottom panels show graphs for relative quantifications of HMT activities in absence (0, MBP) or presence of ULT1 (MBP-ULT1, ULT1), for PRC2^CLF^ and PRC2^SWN^. Error bars correspond to standard deviation for 3 or 2 replicates. Letters indicate significant differences between samples within a graph (Student’s t-test, one sided paired, P-value < 0.05). **(C-D)** Genome-wide influence of combined loss-of-function in ULT1 and CLF or SWN, on H3K27me3 marks in 8 day-old seedlings, as determined by ChIP-seq: WT, *ult1-3*, *clf-28*, *swn-7, ult1-3 clf-28* and *ult1-3 swn-7* (all in Col-0 ecotype). **(C)** Heatmaps showing differences in H3K27me3 abundance in *clf* and *ult1 clf* compared to WT, in regions with more H3K27me3 in *clf* compared to *ult1 clf* (left: *clf* > *ult1 clf,* n= 4279 regions) and in *swn* and *ult1 swn* compared to WT, in regions with more H3K27me3 in *swn* compared to *ult1 swn* (right: *swn* > *ult1 swn,* n= 1001 regions). Heatmaps represent differentially H3K27me3-marked regions, plus 2kb up-and downstream. **(D)** Heatmaps showing differences in H3K27me3 abundance in *ult1*, *swn*, *clf*, *ult1 swn* and *ult1 clf* compared to WT. Areas displayed contain regions with reduced H3K27me3 in *ult1* compared to WT (n= 1438 regions). Heatmaps represent differentially H3K27me3-marked regions, plus 2kb up-and downstream, k-means clustered with k=3. In panels (C) and (D), differences are depicted as log2 fold-change values with respect to WT in heatmap by a color gradient ranging from green for positive values and higher H3K27me3 abundance in the WT to purple for negative values indicating higher mark abundance in the respective mutant. Each line represents one region, all regions are scaled to the same length.

Second, to evaluate *in vivo*, and at the genome-wide scale, the contribution of ULT1 to PRC2^CLF^ and PRC2^SWN^ respective activities, we analysed the effect of ULT1 loss-of-function in each *prc2* mutant background, by ChIP-seq analyses of H3K27me3 marks. For this, we compared H3K27me3 abundance in the *ult1 swn* and *ult1 clf* double mutants to the *swn* and *clf* single mutants, respectively. Based on the results of the HMT activity assays indicating a role for ULT1 in aiding SWN to a higher extent than CLF, we hypothesized that combining the *swn* and *ult1* mutations should not have a major influence on regions with reduced H3K27me3 in the *swn* single mutant, while oppositely, combining the *clf* and *ult1* mutations should more dramatically worsen the H3K27me3 reduction observed in the single *clf* mutant. Indeed, we found 4279 regions with decreased H3K27me3 signal in the *clf ult1* double mutant compared to the single *clf* mutant, and a much smaller number of regions (1001) with significantly reduced H3K27me3 in *swn ult1* compared to *swn*. Moreover, H3K27me3 reduction was much less pronounced at concerned regions in the latter case. The 4279 regions comprise a large fraction with higher H3K27me3 signal at localised areas in the *clf* mutant compared to WT, and this higher signal is reduced to below WT-levels in the *ult1 clf* mutant. This indicates that the absence of *ult1* has a global influence on SWN’s ability to (over-)compensate for CLF’s absence (Fig. 4C). Building on this observation, we sought to refine the resolution of the respective effects of *clf* and *swn* loss-of-function, through clustering of the 1438 regions with less H3K27me3 in the *ult1* mutant. For most of these 1438 regions, the decrease in H3K27me3 observed in the *ult1* single mutant is reduced to noise level in the *ult1 swn* double mutant, indicating that it largely depends on SWN (Fig. 4D). Differently, concerning the contribution of CLF to these 1438 regions repressed by ULT1, the cluster analysis revealed 3 groups: There are regions with complete (Cluster 1) or partial (Cluster 2) enhancement of the H3K27me3 signal in *clf*, and regions with reduced abundance (Cluster 3) in *clf* mutant (Fig. 4D). The enhancement is strongly reduced in the *ult1 clf* double mutant, again hinting at a role for ULT1 in ensuring SWN activity. The fact that regions repressed by ULT1 do not all react similarly to a loss in CLF function indicates distinct relative contributions of the PRC2^CLF^ and PRC2^SWN^ complexes. This difference however is unlikely to reflect distinct targeting, as CLF and SWN were previously reported to sensibly bind to the same chromatin regions (Shu *et al*., 2019). Similar conclusions can be made when analysing ULT1 loss-of-function effect on regions with a H3K27me3 decrease in the *swn* or the *clf* single mutant (Suppl. Fig. S16). Hence, the function of ULT1 as a strong, direct enhancer of PRC2^SWN^ *in vitro*, is concordant with its genome-wide effect observed in mutant combinations with CLF or SWN loss-of-function lines. Overall, these data point ULT1 as a PRC2 cofactor, largely functioning through the enhancement of PRC^SWN^ complex activity.

### Genetic analyses of *ult prc2* mutants support ULT bivalent interactions with CLF and SWN

We previously found that loss-of-function in ULT1 suppressed *clf* phenotypes, leading to qualification of ULT1 as an antagonist of CLF (Carles & Fletcher, 2009). In order to assess the effect of a full loss of the ULTRAPETALA function on the phenotypes induced by a full loss of PRC2 sporophytic function, we combined mutants for ULT1 and its redundant homologue ULT2, with the two HMTs CLF and SWN. We indeed previously reported that both ULT factors play overlapping roles in Arabidopsis development and gene regulation, with ULT1 playing a major role and ULT2 a minor contribution (Monfared *et al*., 2013). The triple *clf-28 swn-7 ult1-3* and quadruple *clf-28 swn-7 ult1-3 ult2-2* mutants were indistinguishable from the double *clf-28 swn-7* mutant. They grow as callus-like tissue with random organs that arise in a disorganized phyllotaxy: ectopic roots, somatic embryos, shoot-like structures (Fig. 5A, B, Suppl. Fig. S17). Hence, loss in ULT function does not recover nor further worsens the highly pleiotropic defects induced by the loss of PRC2 sporophytic function. These results are consistent with our observations indicating that ULT1 promotes the HMT activity of PRC2^SWN^ and, to a lesser extent that of PRC2^CLF^ (Fig. 4). Indeed, if stimulation of PRC2 represents the primary global function of ULT, then no phenotypic rescue would be expected in a *clf swn* double mutant, in which the catalytic subunits of the PRC2 complex are inactive. Accordingly, the *clf swn ult* triple (or quadruple) mutant does not display suppression of the *clf swn* phenotype.

**Figure 5.**
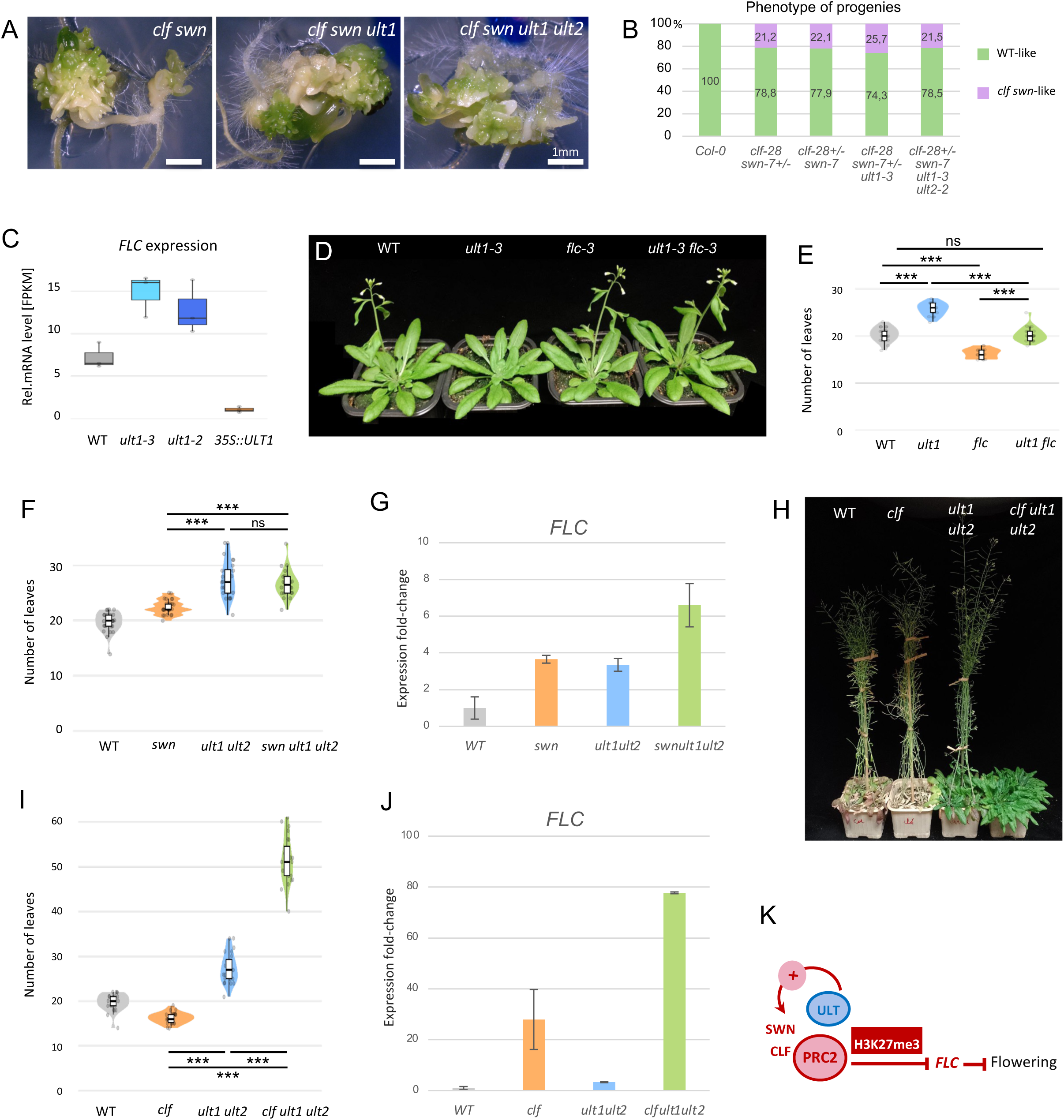
ULT functions in a common pathway with PRC2 for flowering time and *FLC* regulation. **(A,B)** *ult* loss-of-function does not complement sporophytic *prc2* mutant phenotypes. Pictures of double *clf-28 swn-7, triple clf-28 swn-7 ult1-3* and quadruple *clf-28 swn-7 ult1-3 ult2-2* mutants grown for 1 month. All genotypes grow as callus-like tissue with random organs arising in a dis-organized phyllotaxy: ectopic roots, somatic embryos, shoot-like structures. Scale bar: 1mm. Phenotyes of progenies of double *clf-28/+ swn-7, triple clf-28/+ swn-7 ult1-3* and quadruple *clf-28/+ swn-7 ult1-3 ult2-2* mutants, scored 10 days after sowing. Distribution of populations are given in percentages, calculated from 200 plants per progeny. **(C)** Relative transcript levels (FPKM) on *FLC* from RNA-seq data from *ult1-3, ult1-2* mutants and 35S::ULT1 in 8-day-old seedlings grown under LD conditions at 18°C. **(D-E)** Introduction of *flc-3* mutation rescues the late-flowering phenotype of *ult1*. (D) Representative Col-0, *ult1-3*, *flc-3* and *ult1-3 flc-3* mutant strains. (E) Flowering time of Col-0 and *ult1-3*, *flc-3* and *ult1-3 flc-3* mutant strains measured as rosette leaves at bolting. **(F-G)** ULT1 functions in the same pathway as SWN for *FLC* regulation. (F) Flowering time of Col-0, *swn-7, ult1-3 ult2-2* and *swn-7 ult1-3 ult2-2* mutant strains measured as rosette leaves at bolting. (G) Transcript levels of *FLC* measured by qRT-PCR in Col-0, *swn-7, ult1-3 ult2-2* and *swn-7 ult1-3 ult2-2* in 8-day-old seedlings grown under LD conditions at 18°C. **(H-J)** ULT1 functions in the same pathway as CLF for *FLC* regulation. (H) Representative Col-0, *clf-28, ult1-3 ult2-2* and *clf-28 ult1-3 ult2-2* mutant plants grown at LD 20 °C. (I) Flowering time of Col-0, *clf-28, ult1-3 ult2-2* and *clf-28 ult1-3 ult2-2* measured as rosette leaves at bolting. (J) Transcript levels of *FLC* measured by qRT-PCR in Col-0, *clf-28, ult1-3 ult2-2* and *clf-28 ult1-3 ult2-2* in 8-day-old seedlings grown under LD conditions at 18°C. *FLC* expression is strongly upregulated in the triple *clf-28 ult1-3 ult2-2* mutant as compared to the double *ult1-3 ult2-2* mutant (Student’s t-test, one sided paired, P-value< 0.05). In E, F, I, symbols for significant differences correspond to: * P-value < 0.05, ** P-value < 0.01, *** P-value < 0.001; ns: no significant difference (for P-value <0.05). **(K)** Model for *FLC* regulation by ULT1 and PRC2. ULT1 functions together with SWN to promote *FLC* repression, in a pathway synergistic to CLF.

### *FLC* and *WUS* genes are key nodes for ULT1-PRC2 cooperative regulation of developmental transitions

The phenotypes of ULT1 lof (late flowering) and ULT1 gof (early flowering) lines consistently provide evidence that ULT1 functions as a promoter of the reproductive transition (Carles *et al*., 2005); Carles & Fletcher, 2009) (Suppl. Fig. S18). Moreover, the central repressor of flowering *FLC*, master target gene for regulation by PRC2, also stands out as a gene repressed by ULT1 (Tables S1, S3, Suppl. Fig. S18C, Fig. 5C). By generating an *ult1flc* double mutant, in which *flc* lof suppresses *ult1* flowering delay, we revealed that *FLC* is indeed responsible for the late flowering transition in *ult1* mutant (Fig. 5D, E).

Because *FLC* is also a target of SWN and CLF (Shu *et al*., 2020), we investigated the flowering phenotypes of triple mutants lacking both ULT and SWN (or CLF) functions. We could not detect significant differences between triple mutants lacking both ULT and SWN, compared to double mutants lacking ULT function (Fig. 5F). Moreover, *FLC* expression is upregulated in both *ult1 ult2* double and *ult1 ult2 swn* triple mutants, and brought to a yet higher level in the triple mutant (Fig. 5G). This later observation could be explained by a cooperative role of ULT for both sporophytic PRC2 HMT activities. Strikingly, triple mutants lacking both ULT and CLF functions were dramatically delayed in flowering (Fig. 5H, I), with a strong upregulation of *FLC* expression (Fig. 5J). We interpret this as the manifestation of ULT role in strongly enhancing SWN function, which is normally partly masked in the presence of CLF (Fig. 5K), likely because CLF can replace SWN. Furthermore, the weaker late flowering time phenotypes and *FLC* de-repression in *swn ult1 ult2* triple mutant as compared to the *clf ult1 ult2* triple mutant (Fig. 5F) indicates that CLF may compensate better for the absence of SWN than SWN does compensate for the absence of CLF. Hence, the *clf ult1 ult2* plants likely reveal how a *clf swn* double mutant would flower if it could establish correct meristematic structures and body plan. These results also reveal a synergistic interaction between ULT and CLF for *FLC* regulation.

The production of supernumerary floral organs in *ult1* mutant lines was previously attributed to an expansion in the expression domain of the *WUS* meristem regulator (Carles *et al*., 2004, 2005). We found that flowers of triple mutants lacking both ULT and SWN functions produce the same number of organs, for each whorl, as do flowers of double mutants lacking the ULT function only (Suppl. Fig. S19A, B). Differently, triple mutants lacking both ULT and CLF functions produce flowers with more floral organs than *ult1 ult2* double mutants (Suppl. Fig. S19C, D). This indicates a synergistic interaction between ULT and CLF, which could be due to ULT being a strong cofactor of SWN in repressing *WUS*. Moreover, another pathway could be considered for ULT, independent of PRC2, involving a direct interaction with the UIF1 transcription factor which has a repressive function on *WUS* expression (Suppl. Fig. S19E) (Moreau *et al*., 2016).

The similar phenotypes of *ult* and *ult swn* lines, both for flowering time and flower organ numbers, testify for the cooperativity of ULT and SWN. Our genetic analyses, through the two case examples of *FLC* and *WUS*, thus support the model in which ULT1 helps the PRC2 complex to repress genes by enhancing the H3K27me3 histone methyltransferase activity, in particular of PRC2^SWN^; but in the absence of functional SWN, CLF efficiently compensates for H3K27me3 deposition. ULT1 thus likely stands as a moderator of PRC2 function via strong enhancement of SWN activity.

## Discussion

In this study, we report for the first time the genome-wide effect of the ULT1 protein on the abundance of the PcG/PRC2-mediated H3K27me3 and the trxG-mediated H3K4me3 marks, along with its effect on the transcriptome. In order to avoid the consequences of developmental defects but rather observe their epigenetic origin, we ran ChIP-seq and RNA-seq analyses on young seedling tissue (8do). This allowed us to detect significant populations of genes differentially marked with H3K27me3, and slighter effects on H3K4me3 and on gene expression levels. This likely testifies that ULT1 functions post-embryonically as a regulator of PcG/PRC2 activity, rather than as a regulator of trxG activity.

Using AP-MS, Y2H and *in vitro* assays, we demonstrated that ULT1 directly interacts with CLF and SWN within the PRC2 complex. Interaction tests of ULT1 with SWN resulted in faster yeast growth and greater *in vitro* enrichment compared to CLF. By coupling these analyses with HMT assays, we uncovered a novel function for the ULT1 protein as a direct PRC2 activator. These results also highlighted an important difference between the PRC2 complexes carrying either the CLF or the SWN enzyme (PRC2^CLF^ vs PRC2^SWN^). Using recombinant nucleosomes as substrate, the basal activity of PRC2^SWN^ appears to be lower than that of PRC2^CLF^. Importantly, under these experimental conditions, ULT1 markedly enhances the enzymatic activity of PRC2^SWN^ activity, while it has a limited effect on the activity of PRC2^CLF^. This is coherent with observations made using single and double mutants affected in CLF and SWN (Chanvivattana *et al*., 2004). Indeed, while the sporophytic phenotypes of *clf* null mutants are severe (dwarf plants with upward curled leaves and early flowering), those of *swn* null mutants are very subtle (no visible morphological defect, only a slight delay flowering transition). However, the absence of both sporophytic HMTs has dramatic effects, double *clfswn* null mutants being unable to establish a proper plant body plan. This indicates that, while the PRC2^CLF^ complex might be more efficient at repressing common target genes than the PRC2^SWN^ complex, SWN can effectively exert an activity similar to that of CLF in its absence. The two corresponding functions are therefore paralogous with an unequal redundancy. Mutual complementation tests support that CLF and SWN are not interchangeable; indeed, while *clf* plants are complemented by a *35S::CLF* transgene, they are not complemented by a *35S::SWN* transgene (Chanvivattana *et al*., 2004). Interestingly, this is unlikely to be due to different genes bound by CLF or SWN, since they share a large majority of targets, as indicated by ChIP-seq profiling (Shu *et al*., 2019). Based on CLF/SWN overlapping target genes and, on our results, we propose that the difference in gene repression efficiency between PRC2^CLF^ and PRC2^SWN^ is likely due to differential regulation of their enzymatic activity to which ULT1 may contribute. Indeed, our HMT assays reveal that ULT1 selectively enhances PRC2^SWN^ on recombinant substrate and that the presence of post-translational modifications on the nucleosome substrate modulate PRC2^SWN^ and PRC2^CLF^ enzymatic activity.

Nevertheless, an additional and non-exclusive regulatory layer, involving differential binding specificity or affinity to targets, could also explain the differential *clf* vs *swn* defects observed in plants. Further investigations, assessing e.g. the relative affinities of PRC2^CLF^ and PRC2^SWN^ complexes for nucleosomes and chromatin targets, both biochemically and *in planta* will be needed to elucidate the molecular details of this mechanism.

While it would be tempting to compare PRC2^EZH1^/PRC2^EZH2^ to PRC2^SWN^/PRC2^CLF^ according to their respective contributions to H3K27me3 global levels (PRC2^EZH1^ and PRC2^SWN^ being on the lower side), their regulation seems to obey different rules. Indeed, in addition to its lower basal activity, PRC2^EZH1^ tends to respond less efficiently to activating cues such as H3K27me3-mediated allosteric activation (Lee *et al*., 2019). In contrast, we showed that PRC2^SWN^ responds better to ULT1-mediated enzymatic activation, not only *in vitro* but also *in vivo*, where a larger number of genomic regions is impacted by ULT1 loss-of-function, with a markedly stronger reduction in H3K27me3, in the absence of PRC2^CLF^ compared to that of PRC2^SWN^.

Taken together, these results prompt further mechanistic and structural investigation into PRC2^SWN^ and PRC2^CLF^ regulation, in the absence and presence of ULT1.

In plants, the ULT function (ULT1, combined to its functional homologue ULT2 (Carles *et al*., 2005)) is likely responsible for the increase of PRC2^SWN^ activity to the level of PRC2^CLF^ activity. Indeed, we previously reported that the ULT1 and ULT2 proteins are involved in common key developmental processes such as flowering time and organogenesis, through the regulation of inflorescence meristem activity (Carles *et al*., 2005); (Monfared *et al*., 2013). Interestingly, an *emf2 vrn2* double mutant, impaired in two SU(Z)12 alternative PRC2 subunits, when constructed with the weak *emf2-10* allele, displays a late flowering phenotype and additional floral organs (Muller-Xing *et al*., 2014), phenotypes very reminiscent of *ult* mutants. This strongly supports that the severe delay in reproductive transition displayed by the *clf ult1 ult2* triple mutant likely reveals the hidden, joined function of the two alternative CLF and SWN HMTs for *FLC* repression. This function cannot be seen in a *clf swn* double mutant, as it no longer develops as a real plant but a cluster of randomly differentiated cells and tissues. The same is true for the extra flower organs observed in the *clf ult1 ult2* triple mutant, which are significantly more numerous than in *a ult1 ult2* double mutant. Our interpretation of these results is that when CLF is present, repression of *FLC* or *WUS* still occurs, but if CLF is missing, the relay is taken to some extent by SWN, whose function is enhanced by ULT. The *FLC* and *WUS* loci thereby provide demonstrative examples for the developmental effect of ULT as cofactor of PRC2, which moreover discriminates between CLF and SWN. This discrepancy could be explained by ULT1’s role as strong enhancer of PRC2^SWN^, which may in some way compete with the more efficient activity of PRC2^CLF^. The source-sink chromatin model can potentially explain the ULT1/SWN/CLF relationship (Murphy & Berger, 2023). We show that ULT1 increases the H3K27me3 catalytic activity of PRC2^SWN^, and promotes H3K27me3 deposition at a large set of regions at the chromatin. Such a stimulation of PRC2^SWN^ activity (source A) could compete with PRC2^CLF^ activity (source B) via titrating a finite amount of target sites (sinks) to be methylated at H3K27, making ULT1 and SWN apparent repressors of CLF. More generally, in this scenario, two source factors (the H3K27me3 depositing enzymes CLF and SWN), commonly or differentially activated by ULT1, may compete for the same DNA/histone sink locations where stimulation of one source factor would lead to the reduced activity of the other.

Concerning ULT1 de-repressor function, molecular genetic analyses indicate that ULT1 functions independently from the JmJ demethylases. An alternative, remaining possibility is that the reduction in H3K27me3 amounts may rather occur via ULT1 interaction with DNA binding transcription factors (TFs), leading to eviction of PRC2 from target genes, as previously found for AG (Sun *et al*., 2014). ULT1 has been reported to directly interact with TFs, such as members of the MYB and GARP families, in flower meristems (Pires *et al*., 2014; Moreau *et al*., 2016). Through such interactions, it could thus prevent K27 methylation of *de novo* synthesized Histone 3 proteins in dividing stem cells, where it is preferentially expressed, to prompt them for an expression switch at certain genes (Carles *et al*., 2005). However, it is yet unknown whether these ULT1-interacting TFs can oppose the deposition of H3K27 methylation. Consistent as well with a competition scenario against the propagation of the H3K27me3 mark, the ULT1 IP-MS dataset includes potential ULT1 physical partners that either physically or genetically interact with PRC2, or are involved in DNA replication processes: ENHANCER OF LHP1 (EOL1), EARLY IN SHORT DAYS 7 (ESD7) and INCURVATA 2 (ICU2) (Barrero *et al*., 2007; del Olmo *et al*., 2016; Zhou *et al*., 2017; Jiang & Berger, 2017; Godwin & Farrona, 2022). ULT1 may therefore prevent histone mark maintenance during cell division, through interaction with these components.

To date, no robust chromatin profiling dataset has established a clear correlation between ULT1 occupancy and its effects on chromatin marks. Indeed, we and others could not obtain significant overlap between independent ULT1 ChIP-seq datasets (Xu *et al*., 2018; Xu *et al*., 2024), nor between the populations of ULT1-bound genes and ULT1_rep or ULT1_derep genes. While indirect effects cannot be ruled out as a cause for this lack in correlation, it should be noted that the time point of analysis (8do) revealed very little differences in gene expression between the ULT1 lof and WT plants, making indirect effects on a large scale unlikely.

Several technical and biological factors may account for the difficulty in obtaining reliable ULT1 ChIP data. From a technical perspective, steric hindrance arising from ULT1 interactions with chromatin-associated partners, such as the PRC2 complex, may reduce antibody accessibility and compromise immunoprecipitation efficiency. In addition, the presence of an epitope tag may itself interfere with ULT1 function, potentially limiting its recruitment to chromatin and resulting in low enrichment over background in ChIP-seq experiments. Biologically, the lack of detectable enrichment at loci where ULT1 influences H3K27me3 levels may reflect a transient mode of chromatin association. ULT1 may bind chromatin only briefly, dissociating after facilitating PRC2 recruitment or activity. In this context, ChIP-seq for factor profiling provides only a snapshot of protein occupancy, whereas ChIP-seq for H3K27me3 levels represent the cumulative outcome of PRC2 and ULT1 activity over time. Consistent with this interpretation, similar discrepancies between occupancy and functional output have been reported for PRC2 components such as CLF and SWN in plants (Shu *et al*., 2019), as well as in *Drosophila* (Schwartz *et al*., 2006).

Overall, this study demonstrates that ULT1 acts as a developmental switch that can either stimulate or counteract PRC2 repressive function on distinct sets of genes. Moreover, our integrative approach shows that ULT1 can enhance the activity of PRC2 both *in vitro* and *in vivo*. While another family of factors (ALP1, ALP2) were genetically identified as PRC2 antagonists but turned out to be PRC2 components (Liang *et al*., 2015) (Velanis *et al*., 2020), we provide here the first evidence of an initially described trxG protein directly interacting with PRC2 and enhancing its activity. Interestingly, the human PRC2 accessory subunit AEBP2 (which has no homolog in plants), also plays a dual role, with one isoform primarily expressed during early embryogenesis being a cofactor of PRC2, and the other, predominant in adult tissues being an inhibitor of PRC2 (Mucha *et al*., 2025).

JARID2, another human PRC2 accessory subunit can promote PRC2 activity, and PRC2-mediated methylation of JARID2 itself was shown to be important for this function (Sanulli *et al*., 2015). Yet, a recent report suggests that the positive feedback loop regulating hPRC2 enzymatic activity depending on H3K27me3 and JARID2-K116me3 can compete with each other (Agius *et al*., 2026). In contrast, our results indicate that ULT1 can stimulate PRC2 activity despite not being methylated, which indicates a novel mechanism for PRC2 stimulation, yet to be elucidated. Furthermore, our work provides insights into the redundant and distinct functions of CLF and SWN (Shu *et al*., 2020), both at the molecular and developmental levels, and pinpoints ULT1 as a key factor influencing the differing roles of CLF and SWN at cell fate transition genes. We indeed report the first HMT activity tests for SWN and show that PRC2^CLF^ and PRC2^SWN^ complexes have different catalytic efficiencies. This is in agreement with phenotypic studies revealing that, while mutation in *clf* leads to strong phenotypes, mutation in *swn* is indistinguishable from wild type plants (Chanvivattana *et al*., 2004). More broadly, we revealed a molecular mechanism that lies at the crossroads of antagonism and collaboration between PcG/PRC2 and trxG factors.

## Conclusions

Using an integrative approach combining developmental genetics, genomics and biochemistry, we uncovered a unique and novel function that regulates PRC2 HMT activity in plants. We found that ULT1 counteracts deposition of the repressive H3K27me3 mark at hundreds of developmental genes independently of histone demethylases. Additionally, ULT1 promotes PRC2 activity at over a thousand of other developmental genes. Because ULT1 strongly enhances the histone methyl transferase activity of the PRC2^SWN^ complex *in vitro*, and directly interacts with SWN both *in vitro* and *in vivo*, we propose that ULT1 facilitates H3K27me3 deposition at the chromatin, primarily by directly promoting primarily the PRC2^SWN^ enzymatic activity, and to some extent also that of PRC2^CLF^.

Together, our discoveries reveal a mechanism by which developmental transitions at the shoot apex are driven. First, they decouple inhibition of methylation from active demethylation of H3, providing insights into how the switch from H3K27me3-repressed to active genes operates in meristematic regions. Second, they provide a novel mechanism of PRC2 activity enhancement, in which ULT1, via direct interaction, raises deposition of H3K27me3 on nucleosomes, through a mechanism that appears different from that of the PRC2 accessory factors in animals.

## Online Methods

### Arabidopsis mutants and transgenic lines

The *Arabidopsis thaliana* (Arabidopsis) lines used in this study were in the Columbia-0 (Col-0) or Landsberg *erecta* (L*er*) ecotype backgrounds. The mutant lines *ult1-2*, *ult1-3* (SALK_074642C, introgressed or not in Ler), *ult2-2* and *ref6-3* were previously described (Carles *et al*., 2005; Lu *et al*., 2011; Monfared *et al*., 2013). The mutant lines *clf-28* (Salk_139371) (Lopez-Vernaza *et al*., 2012) and *swn-7* (Salk_109121), also known as *swn-4* (Wang *et al*., 2006) are in the Col-0 background. The *clf-28/+ swn-7* line was kindly provided by Daniel Schubert and used to cross to the double mutant *ult1-3 ult2-2* in Col-0 (Monfared *et al*., 2013) in order to obtain the quadruple mutant *clf-28 swn-7 ult1-3 ult2-2*.

The constructs used for transgenic lines were introduced in Arabidopsis plants by Floral dip agrobacterium-mediated transformation. *Agrobacterium tumefaciens*, strain C58C1pMP90, carrying the *p35S::ULT1* (Carles & Fletcher, 2009) or the p*35S::REF6-YFP-HA*, (Lu *et al*., 2011) construct were grown overnight at 28°C in 2ml of selective LB medium and 1 ml of this culture was then spread on YEB medium supplemented with antibiotics and grown at 28°C during 72hr. Grown colonies were dissolved in 50ml LB and grown at 28°C during 2hr, before being pelleted and resuspended with 200ml of a 5% sucrose solution containing 0.01% of Silwet L-77. Inflorescences of *ref6-3* or *ult1-3* mutants or Col-0 WT control plants were dipped into this bacterial culture during 10s. The dipped plants (T0) were grown within a controlled environment until their seeds were fully mature. T1 generation seeds were collected from T0 plants and further selected for antibiotic resistance on MS selective medium.

When cultivated *in vitro*, seeds were sterilized (70% ethanol solution for 5 min and 95% ethanol solution for 5 min, dried on Whatman paper), sown on ½ MS medium supplemented with 1% sucrose, and then stratified for 3 days at 4°C. Seeds were further germinated within a controlled environment chamber at 20°C with 16hr light/8hr dark photoperiod and under 65% humidity.

When directly sown on soil, seeds were stratified for 3 days at 4°C, and then transferred in a controlled environment chamber at 20°C with 16hr light/ 8hr dark photoperiod and under 65% humidity.

### Chromatin immunoprecipitation followed by deep sequencing (ChIP-seq)

ChIP-seq was performed on 8-day old seedlings of L*er* WT, *ult1-2*, *ult1-3*, *35S::ULT1*, Col-0 WT and *ref6-3* as described in (Engelhorn *et al*., 2017b); and on 8-day old seedlings of Col-0 WT, *ult1-3*, *clf-28, swn-7, ult1-3 clf-28* and *ult1-3 swn-7*, with chromatin preparation and immuno-precipitation as described in (Zhu *et al*., New Phytol, 2024). Seedlings were grown in long day (LD) conditions of 16 h light (18 degrees) and 8 h darkness (16 degrees) on Murashige and Skoog medium (Sigma-Aldrich, Lyon, France) supplemented with 0.3% sucrose in growth cabinets (Percival, Percival Scientific Incorporated, provided by CLF Plant Climatics, Emersacker, Germany).

Antibodies and quality control were described previously (Engelhorn *et al*., 2017a). Sequencing was performed as described for replicate 2 (Engelhorn *et al*., 2017a) in 50bp single read mode for L*er* WT, *ult1-2*, *ult1-3*, *35S::ULT1*, Col-0 WT and *ref6-3* and on a DNBseq^TM^-G400 platform at BGI (China) in 100 PE mode for Col-0 WT, *ult1-3*, *clf-28, swn-7, ult1-3 clf-28* and *ult1-3 swn-7*.

Read mapping, processing and determination of differentially marked regions for single read data was performed as described in (Engelhorn *et al*., 2017b): in brief, reads were mapped to the Arabidopsis Col-0 background (Tair10) with bwa (Li and Durbin, 2009). The resulting sam files were converted to bam files using samtools. Duplicated reads were removed with Picardtools MarkDuplicates (http://broadinstitute.github.io/picard/). Uniquely mapped reads were employed for peak calling to determine H3K27me3 and H3K4me3 marked regions with significant enrichment (p-value < 0.0001) over input using SICER (Zang *et al*., 2009). Paired-end read data were treated with the same pipeline but adapted to accommodate paired end data: ChIP and input samples with more than 30% higher read numbers than the replicates average were down-sampled to fall within the average (this concerned the input *ult1* and IP WT for Rep.1) using samtools. Bam files were converted into BEDPE format using bedtools (Quinlan and Hall, 2010) (v2.30.0). Paired reads were merged into one fragment using awk (4.1.4). Sub-nucleosomal fragments (<146 bases), large undigested fragments (>500bp), reads mapping to regions prone for artificial signals (Klasfeld *et al*., 2022), a region surrounding the centromere of chromosome 4 (low signal in input of *clf-28*, putative inversion caused by T-DNA) and all reads mapping to the plastid or mitochondria genome were removed at this step. To avoid normalisation problems due to differences in fragment size, all fragments were set to a length of 146 bases around the fragment centre and reads of both replicates were merged for further analysis. In all cases, differentially marked regions (p-value < 0.0001) were determined with the SICER-df function modified to employ a mappable/effective genome size of 0.9. For paired end data, the function was further modified so that the shifting of reads by 0.5x fragment size was omitted. Target genes/ differentially marked genes were determined as genes overlapping with target regions / differentially marked regions with at least 1bp of their gene body, respectively. For replicate 2 of H3K27me3 in *ult1-2* and *ult1-3*, read numbers were so high that peaks displayed saturation. Therefore, for the quantitative comparison, duplicates were not removed. Only regions/genes differentially marked in both replicates were considered for further analysis. To obtain high confidence gene lists, differentially H3K27me3 marked genes with simultaneous lower H3K27me3 signal in both *ult1* lof mutants and higher H3K27me3 signal in the gof line (ULT1_rep), or higher H3K27me3 signal in both *ult1* lof mutants and lower H3K27me3 signal in the gof line (ULT1_derep) were determined by intersection of the differentially marked gene lists.

For comparisons of H3K27me3 differences and patterns in PcG/trxG mutants *clf, swn* and *elf6-3 ref6C jmj13G*, raw data from ChIP-seq experiments performed in (Shu *et al*., 2019) and (Yan *et al*., 2018) was downloaded from GeoAccession GSE108960 and GSE106942 and analysed using the same parameters employed for the libraries generated in this study. In this analysis, we aimed for a stringent p-value cut-off (p-value < 0.0001) instead of a fold-change cut-off to enable detection of subtle changes in H3K27me3 levels over longer stretches as we observed for our ULT1 analysis. It should be noted that due to this analysis method, numbers of genes displaying lower or higher H3K27me3 levels in the different genotypes might differ from previously published results. For *elf6-3 ref6C jmj13G* no input was available, thus an E-value cutoff of 100 was chosen in the SICER analysis (Zang *et al*., 2009).

Significant over- and under-enrichment for comparisons of differentially histone marking target gene overlaps was calculated by hypergeometric testing in R (Version 4.1.2) using the phyper function and all WT Ler H3K27me3 targets as total number of targets.

Normalised bigwig files containing H3K4me3 and H3K27me3 coverage per million reads were generated as described in (Engelhorn *et al*., 2017b) using bash/awk and bedGraphToBigWig (Kent et al. 2010). Heatmaps, including generation of log2-fold change bigwig files for comparisons and metagene plots were produced from these bigwig files with deeptools (v3.5.1) (Ramírez *et al*., 2016). Genomic coverage screenshots were taken with Integrated Genome Browser (v10.0.0) (Freese *et al*., 2016) and the Integrative Genomics Viewer (IGV, v.2.6.3)(Robinson *et al*., 2011).

### ChIP-qPCR

For ChIP-qPCR independent IPs were performed on the same material as employed for ChIP-seq plus a third replicate. The same protocol was followed for the ChIP-seq IPs with the exception of the sonication. In order to obtain longer fragments for qPCR, sonication was performed for 10 min with 15 sec on 1 min off cycles in a Bioruptor machine (Agilent) filled with ice-cold water at strength H. Nuclear extracts were diluted 1:10 and 2 ul of this dilution were used as template in a total reaction volume of 10 ul using SsoFast Eva Green Supermix with primers at 0.4 uM (primer sequences are provided in Table S6). Technical triplicates were performed for each sample analysed. The quantitative PCR reaction was performed on a Biorad CFX Connect machine.

### RNA extraction followed by deep sequencing (RNA-seq)

RNA-extraction, sequencing and analyses were performed on RNA prepared from the same tissues as those used for chromatin preparations as well as a third replicate, as described in (Engelhorn *et al*., 2017a) and (Engelhorn *et al*., 2017b), using TopHat v2.1.1, cuffdiff v2.2.1. For comparison of single gene counts, Fragments per kilobase per million read (FPKM) values were generated using cuffnorm v2.2.1 with library-norm-method geometric (Trapnell *et al*., 2012). For ULT1 gof, an independent insertion was chosen as the third replicate. All of these *35S::ULT1* replicates show ULT1 over-expression in RNA-seq (ULT1 FPKM: 685.3 809.8 and 512.4 in rep1-3; Ler FPKM: 16.8, 10.2 and 10.3 in rep1-3). Volcano plots were generated with ggplot2 (v3.4.4) [https://ggplot2.tidyverse.org].

### Gene Ontology analyses

GO enrichment analyses were performed in different gene sets using ShinyGO tool v0.8 (Ge *et al*., 2020). We used the WT Ler marked genes for either H3K27me3 or H3K4me3 as a background. Terms were sorted by increasing −10log(FDR).

### Western-blots on marks

600mg of 8-day-old seedlings were frozen and ground in liquid nitrogen to a fine powder and homogenized in Honda’s buffer. Nuclei were isolated as previously described (Xia *et al*., 1997). Nuclear pellets were resuspended in 2X Laemmli buffer, supplemented with β-mercaptoethanol (Bio-Rad). The volume of Laemmli buffer was adjusted to the size of the protein pellet obtained after extraction. Protein concentration was determined in each sample using the Qubit Protein Assay Kit (Invitrogen) and a first blot was run to assess for even loading, using an anti-H3 antibody. After signal quantification, the volumes of the extracts were eventually adjusted for optimal loading before performing detection and quantification of H3K27me3, H3 and H4. After incubation for 5 min at 90°C and centrifugation at 16000 g, nuclear protein extracts were loaded on 4-20% precast SDS-PAGE gels (Bio-Rad) for Western Blot analysis. The primary antibodies were Abcam ab1791 (H3, 1/25000), Diagenode C154110069 (H3K27me3, 1/1000), Millipore 04-858 (H4, 1/5000). The secondary antibody was the horseradish peroxidase-coupled anti-rabbit antibody (Abliance, 1/5000). Detection was performed using BioRad Clarity™ ECL substrate. Densitometric analysis of immunoreactive protein bands was performed on non-saturated signals, using Image Lab software (Bio-Rad, France). The quantified H3K27me3 signal was normalized to H3 or H4 or total protein amount in the sample.

### Immuno-fluorescence and DAPI staining of nuclei

Eight-day-old plants were harvested and fixed for 30 minutes in 4% formaldehyde in the PME buffer (50 mM PIPES, 5 mM MgSO4, 1 mM EGTA, pH 6.5). Samples were washed and transferred to a Petri dish, chopped in enzyme mix (1% cellulase, 0.5% pectolyase, and 1% cytohelicase) and incubated 30 minutes at RT. Suspension was filtered through nylon mesh (pore size 30 um) and spread on poly-lysine coated slides. 20uL of 1% lipsol in PME was added on the slide and spread, then 20uL of 4% paraformaldehyde in PME was added and spread, and the slides left to air dry. Slides were incubated with 0.5% Triton X-100 in 1X PBS for 15 minutes at RT in a moist chamber, washed in 1X PBS, and incubated overnight at 4°C with an anti-H3K27me3 antibody (C154110069, Diagenode, batch A1818P, 1:400) in 3% BSA, 0.05% Tween-20 in 1X PBS then washed and incubated 3 hours at RT with an anti-rabbit antibody coupled to Alexa Fluor™ 488 (A-11070, 1:1000) diluted in 3% BSA, 0.05% Tween-20 in 1X PBS. Slides were mounted in Vectashield containing DAPI and nuclei imaged with a Zeiss LSM 800 AiryScan confocal microscope using a 63X objective.

### RT-qPCR

To quantifiy transcript levels, total RNAs were isolated from 8-day-old seedlings using the RNeasy™ plant mini kit (Qiagen). Reverse transcription was then performed on 1 to 2 ug of total RNA using the Master Mix Superscript™ IV Vilo™ (ThermoFisher Scientific). Quantitative PCR reactions were performed with the Power SYBR™ Green master mix (ThermoFisher Scientific), with forward and reverse primers each at 0.2 uM, in a total reaction volume of 10 ul. The list of primers is provided in primer sequences in Table S6. Technical triplicates were performed for each sample analysed. The quantitative PCR reaction was performed on a Biorad CFX Connect machine. The results were analysed using the REST spreadsheet (Pfaffl *et al*., 2002).

### Affinity Purification followed by Mass Spectrometry (AP-MS)

For affinity-purification coupled to mass spectrometry (AP-MS) experiments, ULT1 coding sequence was cloned in the pCambia2300-5’GStag- polyA vector, at the KpnI and EcoRI sites, to generate a GS tag-ULT1 fusion (tag in the N-terminus end of ULT1), expressed under the control of the CaMV35 promoter. Arabidopsis suspension cultured PSB-D cells were maintained, transformed and selected as previously described (Van Leene *et al*., 2015). Affinity purification was performed as described in (Pfab *et al*., 2017). First, rabbit IgG (Sigma) was coupled to epoxy-activated magnetic beads (Bioclone). Buffer composition markedly influences the integrity of protein complexes isolated by affinity purification. Therefore, mild conditions (25 mM HEPES-KOH pH 7.4, 0.05% IGEPAL CA-630, 1 mM DTT, 100 mM NaCl, 2 mM MgCl2, 5 mM EGTA, 10% glycerol, cOmplete™ EDTA free proteinase inhibitor tablets (Sigma-Aldrich), 1 mM PMSF for extraction/washing buffer) were utilized for IgG affinity purification. Benzonase-treated protein extracts of 15g of *A. thaliana* PSB-D suspension cultured cells (Van Leene et al., 2015) were used to isolate GS-tagged ULT1 and co-purifying proteins.

Protein samples eluted from the IgG-beads were were separated using NuPAGE 4-12 % SDS gels (Thermo Scientific, Dreieich). After staining and destaining each lane was cut into slices, transferred into 2 ml micro tubes (Eppendorf) and washed with 50 mM NH_4_HCO_3_, 50 mM NH_4_HCO_3_/acetonitrile (3+1) and 50 mM NH_4_HCO_3_/acetonitrile (1/1) while shaking gently in an orbital shaker (VXR basic Vibrax, IKA). Gel pieces were lyophilized after shrinking by 100 % acetonitrile. To block cysteines, reduction with DTT was carried out for 30 min at 57°C followed by an alkylation step with iodoacetamide for 30 min at room temperature in the dark. Subsequently, gel slices were washed and lyophilized again as described above. Proteins were subjected to in gel tryptic digest overnight at 37°C with approximately 0.5 μg trypsin (Trypsin Gold, mass spectrometry grade, Promega). Peptides were eluted twice with 100 mM NH_4_HCO_3_ followed by an additional extraction with 50 mM NH_4_HCO_3_ in 50 % acetonitrile. Prior to LC-MS/MS analysis, combined eluates were lyophilized and reconstituted in 20 μl of 0.1 % formic acid. Separation of peptides by reversed-phase chromatography was carried out by an UltiMate 3000 RSLCnano System (Thermo Scientific, Dreieich) which was equipped with a C18 Acclaim Pepmap100 preconcentration column (100 μm i.D. x 20 mm, Thermo Fisher) in front of an Acclaim Pepmap100 C18 nano column (75 μm i.D. x 250 mm, Thermo Fisher). A linear gradient of 4 % to 40 % acetonitrile in 0.1 % formic acid over 60 min was used to separate peptides at a flow rate of 300 nl/min. The LC-system was coupled on-line to a maXis plus UHR-QTOF System (Bruker Daltonics, Bremen) via a CaptiveSpray nanoflow electrospray source (Bruker Daltonics). Data-dependent acquisition of MS/MS spectra by CID fragmentation was performed at a resolution of minimum 60000 for MS and MS/MS scans, respectively. The MS spectra rate of the precursor scan was 3.5 Hz processing a mass range between m/z 50 and m/z 2200. Via the Compass 1.7 acquisition and processing software (Bruker Daltonics) a dynamic method with a fixed cycle time of 3 s and a m/z dependent collision energy adjustment was applied. Protein Scape 3.1.3 (Bruker Daltonics) in connection with Mascot 2.5.1.1 (Matrix Science) facilitated database searching of the TAIR10 database. Search parameters were as follows: trypsin/P, 2 missed cleavages allowed, deamidation (N, Q), oxidation (M), carbamidomethyl (C), propionamide (C) and acetylation (protein N-terminal) as variable modifications, precursor tolerance 0.02 Da, MS/MS tolerance 0.04 Da, significance threshold p<0.05. Mascot peptide ion-score cut-off was set to 15. A protein score of minimum 30 and at least 2 peptides found with an individual ion-score of ≥ 15 were considered as criteria for reliable protein identification.

### Yeast-two-hybrid (Y2H) assays

For binary interaction tests, the candidate partners were expressed as Gal4DNA-BD (BD: binding domain) or Gal4AD (AD: activation domain) fusion proteins. For this, all cDNAs were cloned into pGAD-T7 and pGKB-T7 vectors (Clontech) and were transformed into yeast strains AH109 and Y187, respectively, for mating according to the manufacturer. ULT1 and UIF1 full-length and truncated versions were previously described in (Moreau *et al*., 2016) and CLF and SWN clones were previously described in (Chanvivattana *et al*., 2004). A shortened version of SWN, SWNΔSET (SWN_1-696_) in which the histone methyl transferase SET domain is absent, was used in these experiments. ULT1 and CLF or SWN-transformed yeast strains were mated and the resulting diploids were selected for the presence of both plasmids with medium lacking Leu and Trp (−LW). Selected yeast strains (containing both a pGADT7 vector and a pGBKT7 vector) were resuspended in water at two densities, OD600nm=10 and OD_600nm_=1. Drops of 5uL of resuspended yeast were then dotted on −LW or –LWAH medium (lacking Leu, Tryp, His and Ade) on which only the diploids presenting protein interactions can grow. The plates were incubated at 30°C for 3 to 5 days and yeast growth was monitored every day.

### Production and purification of recombinant PRC2 and ULT1

To produce the catalytic lobes of the two PRC2 complexes in the baculovirus expression system, His-tagged full-length CLF and SWN were respectively cloned into the Multibac acceptor vector pFL (Fitzgerald *et al*., 2006). Strep-tagged full-length FIE and C-terminally Flag-tagged EMF2 VEFS domain (residues 468-631) were respectively cloned into the donor vectors pUCDM and pIDK. The two donor vectors were recombined with the acceptor pFL carrying either CLF or SWN using Cre-recombination, generating two Multibac plasmids coding for either trimeric PRC2^SWN^ or PRC^CLF^. The PRC2 plasmids were inserted into the bacmids DNA and used to transfect Sf21 insect cells, resulting in the PRC2-producing baculoviruses. The complexes were expressed by infecting High Five cells with the baculoviruses at 27°C for 72 hours. Following cell disruption, supernatants containing soluble complexes were incubate for 2 hours with Ni^2+^-Chelating Sepharose (Cytiva). The resin was washed with a buffer containing 25 mM HEPES, pH 7.8, 150 mM NaCl, 1 mM TCEP, 2 mM MgCl_2_, and 10 mM imidazole, followed by a high-salt buffer wash with 1 M NaCl. The bound complex was eluted by increasing the imidazole concentration to 300 mM. To obtain a stoichiometric complex, the eluted complex was further purified by Strep-Tactin XT resin (IBA) and size exclusion chromatography on a Superdex 200 Increase (Cytiva) in a buffer containing 25 mM HEPES pH 7.8, 150 mM NaCl, 1 mM TCEP, 2 mM MgCl2 and 2% glycerol.

Full-length ULT1 was expressed in E. coli Rosetta 2 cells (Novagen) from the pETM41 vector (EMBL, Gunter Stier) as His-MBP-tag fusion. Before induction, the growth medium was supplemented with 1mM ZnSO_4_. The protein was first purified on Amylose resin (NEB) and the aggregated species were eliminated by gel filtration on Superdex 200 Increase (Cytiva) in a buffer containing 25 mM HEPES pH 7.8, 300 mM NaCl, and 10 mM DTT. After the His-tag removal by TEV protease the protein was further purified on a Heparin HiTrap column (Cytiva) and another gel filtration on Superdex 200 Increase (Cytiva).

### Pull-down assays

The purified PRC2 complexes (PRC2^CLF^ and PRC2^SWN^) and untagged ULT1 protein were prepared in a buffer containing 25 mM HEPES, pH 7.8, 100 mM NaCl, 1 mM TCEP, 2 mM MgCl_2_ and 2% glycerol and mixed in a molar ratio of 1:3, respectively. The mixture was incubated for 30 minutes before being loaded onto Strep-Tactin XT (IBA) resin columns. The resin was extensively washed with a buffer containing 25 mM HEPES pH 7.8, 100 mM NaCl, 1 mM TCEP, 2% glycerol, and 2 mM MgCl_2_, and the bound proteins were eluted by adding 50 mM D-Biotin and analyzed by 15% SDS-PAGE.

### HMT activity assays

HMT assays were performed with 1.5ug of PRC2-CLF or PCR2-SWN alone or in presence of 0.75ug of MBP or MPB-ULT1 or ULT1 incubated at 30°c for 30min in the reaction buffer containing 50 mM Tris-HCl (pH 8.5), 5 mM MgCl2, 4 mM DTT, and 1 uM 3H-labeled SAM (Amersham Pharmacia Biotech). HMT assays were performed with oligo-nucleosomes reconstituted with the four histones from *Xenopus laevis* expressed in *Escherichia coli* (Luger *et al*., 1997). 0.75ug of recombinant histone oligo-nucleosomes were used as substrate. The total volume of the reaction mixture was adjusted to 25 ul. The reaction was stopped by addition of SDS sample buffer and boiled for 5mn at 95°c. The samples were then fractionated by 15% SDS-PAGE, transferred to a PVDF membrane, and stained with Coomassie. The membranes were exposed overnight on a phosphorimager screen and revealed on an imager typhoon (Cytiva).

### Obtention of the clf swn ult1 ult2 multiple mutants

Multiple mutants were obtained from an initial cross between *clf-28/+ swn-7* and *ult1-3 ult2-2* lines, both in the Col-0 ecotype.

#### clf-28 swn-7 ult1-3 triple mutant

The progeny of a *clf-28 swn-7+/- ult1-3* line was grown on ½ MS medium (1% sucrose) for 15 days at 20°C in long days (16hr light/8hr dark). Out of 150 seedlings, 35 displayed a *clf swn* double mutant phenotype and were confirmed to be homozygous for the *swn-7* mutation by genotyping (see primer sequences in Table S6).

#### clf-28 swn-7 ult1-3 ult2-2 quadruple mutant

Due to genetic linkage of approx. 2-4 cM between the *CLF* and *ULT2* loci, we attempted to obtain the *clf-28 swn-7 ult1-3 ult2-2* quadruple mutant first using the CRISPR Cas9 method to create an *ult2* mutant, which turned out unsuccessful, second by a high throughput screen through a population obtained by genetic crossing. For this, we screened through the progeny of a *clf-28/+ swn-7 ult1-3 ult2-2/+* plant. 550 seeds were grown on soil and genotyped for *ult2-2* and *clf-28/+* mutations (primer sequences provided in Table S6). One plant displayed the expected genotype *clf-28/+ swn-7 ult1-3 ult2-2*. Phenotypic observations on the progeny of this plant were carried out *in vitro*.

### Phenotypic analyses of the clf swn ult1 triple and clf swn ult1 ult2 quadruple mutants

200 seeds were sown for each the following genotypes, and grown on ½ MS (1% sucrose), at 20°C (16hr light/8hr dark; 65% humidity): *clf-28 swn-7+/- ult1-3* and *clf-28+/- swn-7 ult1-3 ult2-2* as well as the control lines *clf-28+/- swn-7* and *clf-28 swn-7*. After 15 days of growth, *clf*-like seedlings were counted under the binocular microscope SZX12 and genotyped for *clf-28*, *swn-7* and *ult2-2* mutations.

### Imaging by multifocal microscope and Environmental Scanning Electron Microscope (SEM)

We followed the evolution of the *clf*-like phenotypes after two months of *in vitro* growth, and transplanted the corresponding mutants onto fresh ½ MS medium (1% sucrose) after 1 month. Pictures of these mutants were taken with the VHX-5000 Keyence numeric multifocal microscope as well as with an environmental SEM. SEM experiments were performed at the Electron Microscopy facility of the ICMG Nanobio-Chemistry Platform (Grenoble, France). Untreated tissues were directly placed in the microscope chamber. Care was taken to maintain some humidity during the pressure decrease in the chamber in order to prevent tissue drying. Secondary electron images were recorded with a Quanta FEG 250 (FEI) microscope while maintaining the tissue at 2◦C, under a pressure of 500 Pa and a 70% relative humidity. The accelerating voltage was 14 kV and the image magnification ranged from 100 to 800x.

### Flowering time and Floral organ counting

The flowering times were measured by the leaf counts at bolting. For floral organ counting the first 10 open flowers from 10 plants were counted from each genotype. A Wilcox statistical test was performed on experimental data.

## Acknowledgements

We thank Evelyna Damyanova for help with technical realization of the ChIP used in qPCR, Enora Fremy for preliminary Y2H assays, Renaud Dumas and Catarina Da Silva for help in designing ULT1 protein production strategies, François Parcy for discussions and infrastructure share, Stéphane Ravanel for preliminary HMT assays, Franziska Turck for sharing *clf/swn* ChIP-seq data prior to publication and Christine Lancelon-Pin for SEM manipulations (at Cermav Grenoble), and the iGReD CLIC microscopy facility for image acquisition.

## Author contributions

Conceptualization, C.C.C. and J.E.; methodology, C.C.C, J.E., G.V., J-B. I., J. K. and V.G.; formal analysis, C.C.C., J.E., G.V., J.K., V.G, R. M. and L.T.; investigation, J.E., V.G., C.T., J-B. I., H.L., M.W., M.V., P.M-H., K.D.G., E.T., A.B., J.K., A.P., M.L.M. and C.C.C.; writing – original draft, C.C.C.; writing – review & editing, C.C.C. with help of J.E., J.K., R.M., V.G, M.W., G.V., L.T. and A.P.; funding acquisition for this specific study, C.C.C. and J.K.

---

**Suppl. Table S1.** Lists of ULT1_derep and ULT1_rep target genes. Four sheets are displayed in the file, which correspond to, in order: all 916 ULT1_derep genes; subset of ULT1_derep genes that encode meristem, flowering and flower morphogenesis regulators; all 1362 ULT1_rep genes; subset of ULT1_rep genes that encode meristem, flowering and flower morphogenesis regulators. *See excel File*

**Suppl. Figure S1.**
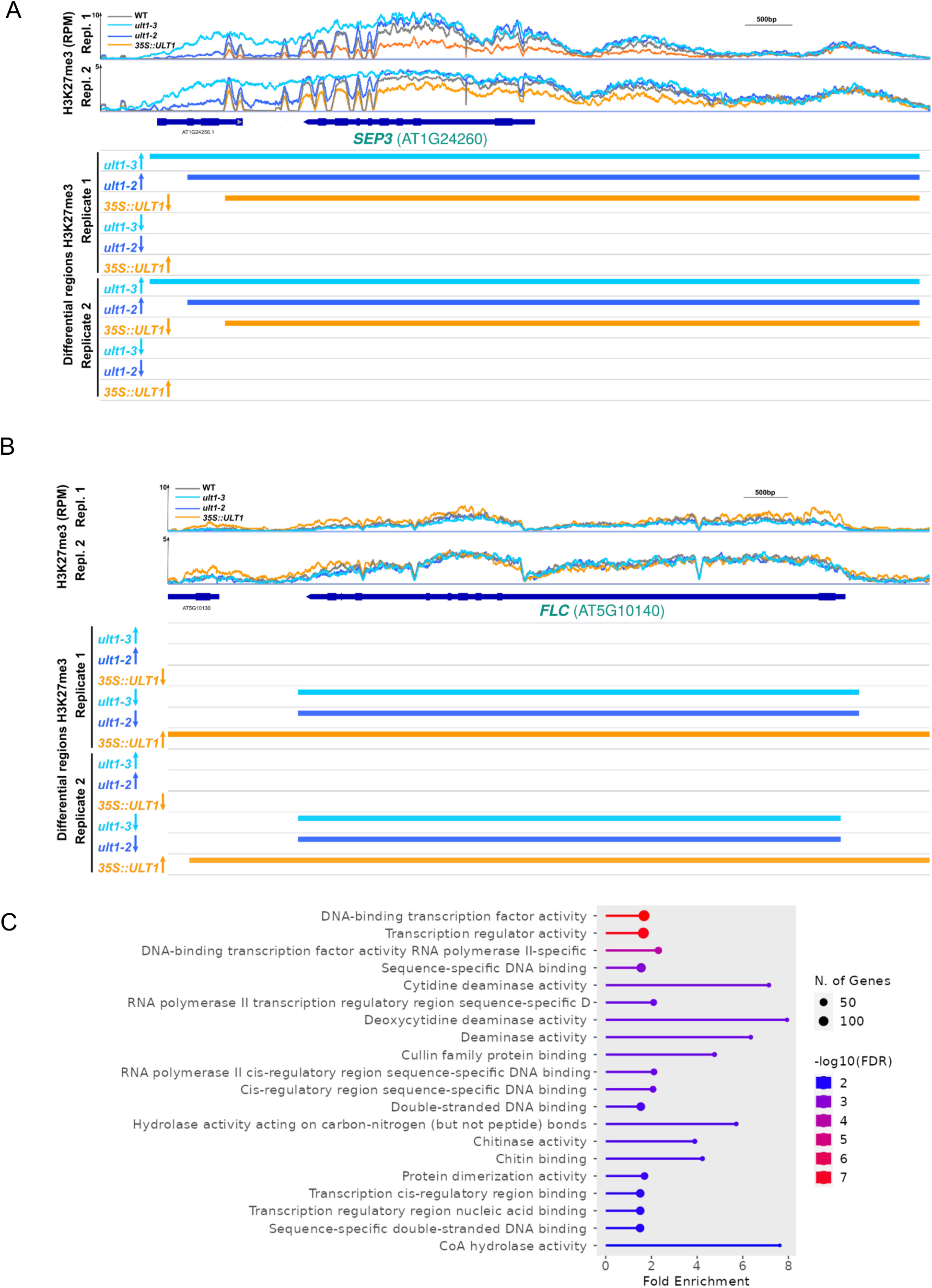
(A-B). Example of SICER-called regions, for differential abundance of H3K27me3 between genotypes. Genome browser (IGV) views of H3K27me3 coverage determined by ChIP-seq (two replicates) in wild type, *ult1-2*, *ult1-3* and *35S::ULT1* lines at (A) *SEPALLATA3* (*SEP3*) (an ULT_derep target) and at (B) *FLOWERING LOCUS C* (*FLC*) (an ULT_rep target). The bars below depict the regions which are called for significantly different abundance of H3K27me3. RPM: reads per million. **(C) Functional categorisation of ULT1_derep genes.** ShinyGO v0.8 analysis for significantly over-represented Gene Ontology terms among ULT1_derep genes compared to H3K27me3 target genes in Ler WT.

**Suppl. Figure S2.**
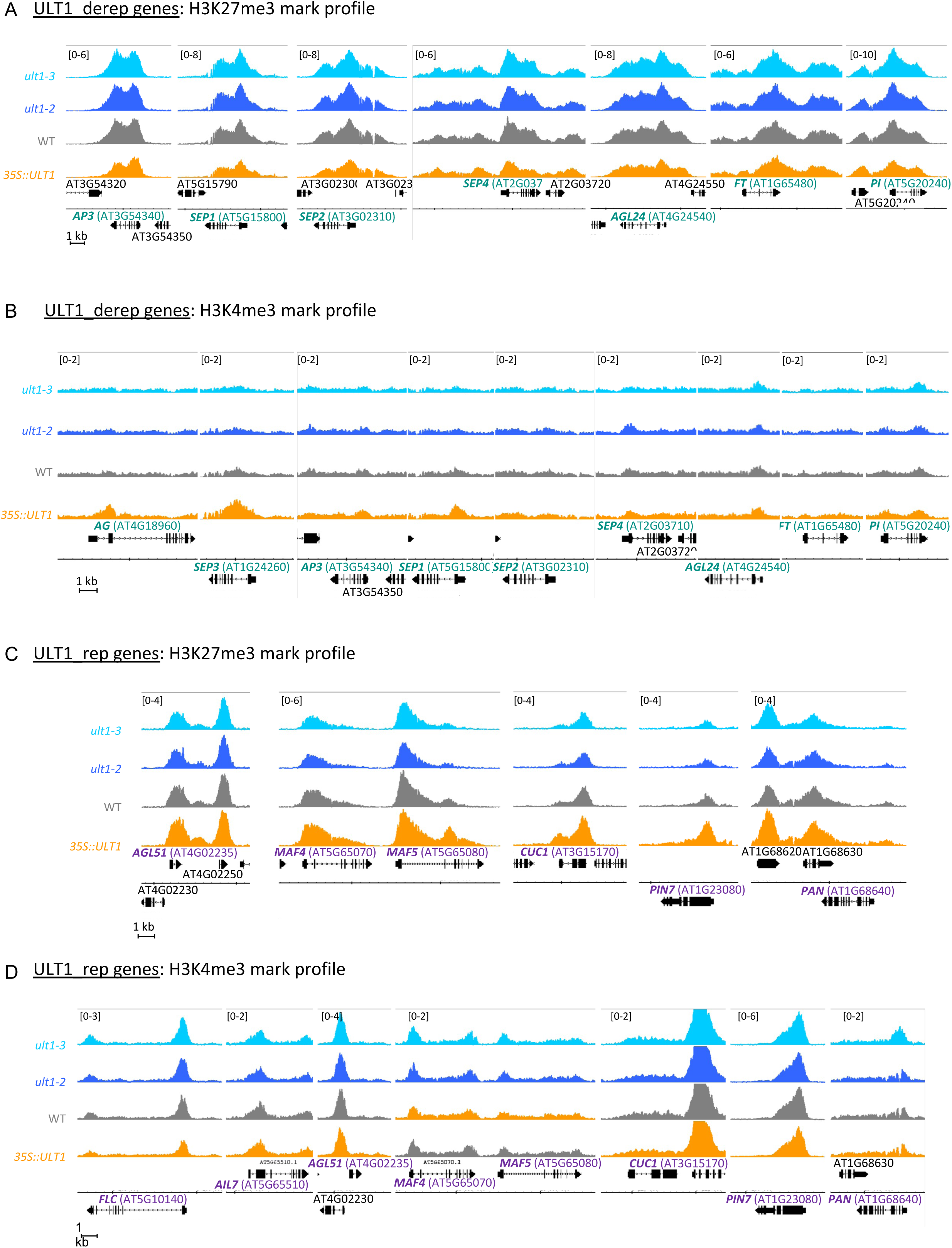
Examples of influence of ULT1 on histone marks. Genome browser (IGB) views of (A,C) H3K27me3 and (B,D) H3K4me3 mark profiles, as determined by ChIP-seq in wild type (Ler), *ult1-2* (EMS line, null mutant), *ult1-3* (T-DNA insertion line, null mutant) and *35S::ULT1*. Data is shown for (A-B) the ULT1_derep flower morphogenesis genes *APETALA3* (*AP3*), *SEPALLATA 1-4* (*SEP1,2,4*), *AP1, PISTILLATA* (*PI*) and the flowering regulators *AGAMOUS-LIKE 24* (*AGL24*) and *FLOWERING LOCUS T* (*FT*), and for the ULT1_rep genes *AGAMOUS-LIKE LOCUS 51* (*AGL51*), *FLOWERING LOCUS C* (*FLC*), master regulator of the reproductive transition and its paralogues MADS AFFECTING FLOWERING 4 (MAF4) and MAF5, the meristem regulators *AINTEGUMENTA-LIKE 7* (*AIL7*), CUP-SHAPED COTYLEDON1 (CUC1), ARABIDOPSIS PIN-FORMED 7 (PIN7) and PERIANTHIA (PAN); unless already shown in Figure 1 for H3K27me3.

**Suppl. Figure S3.**
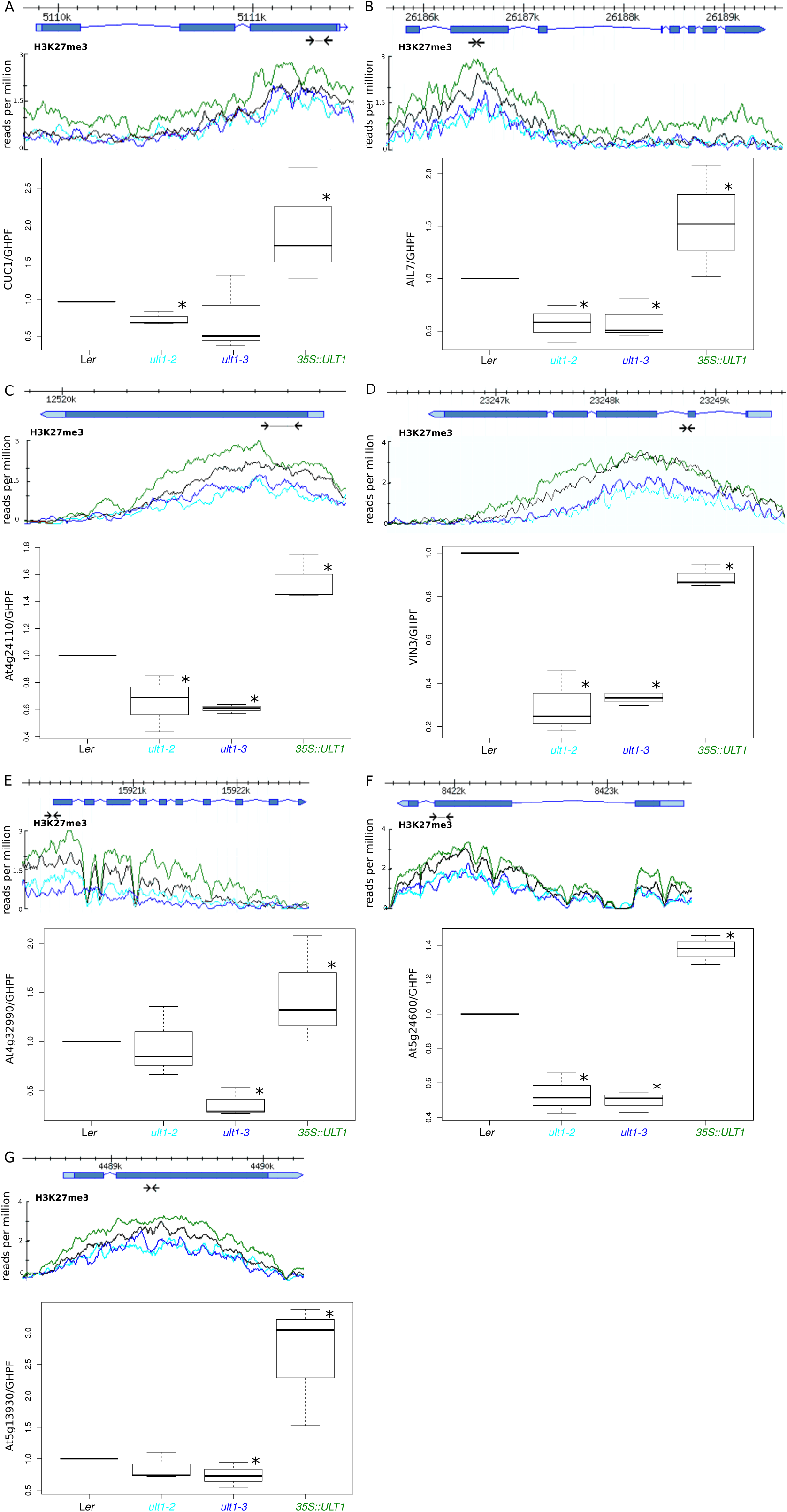
ChIP q-PCR analysis of H3K27me3 abundance in *ult1-2, ult1-3* and *35S::UL*T1 at seven selected ULT1_rep genes. *AT3G62710* (Glycosyl hydrolase family protein - GHFP) was used to normalise the ChIP signal. Box plots show combined data of three biological replicates, the horizontal line displays the median, the whiskers mark the upper and lower values, the box represents the 25 % above and below the median. * reflects to statistical significance with Kruskal-Wallis test followed by post-hoc non-parametric two-sided Tuckey test, using the R package nparcomp (Knell, 2013). For comparison, the ChIP-seq results over each locus are included and the employed primers are indicated by arrows below the gene model. A: *CUC1*, B: *AIL7*, C: At4g24110, D: *VIN3*, E: At4g32990, F: At5g24600, G: At5g13930.

**Suppl. Figure S4.**
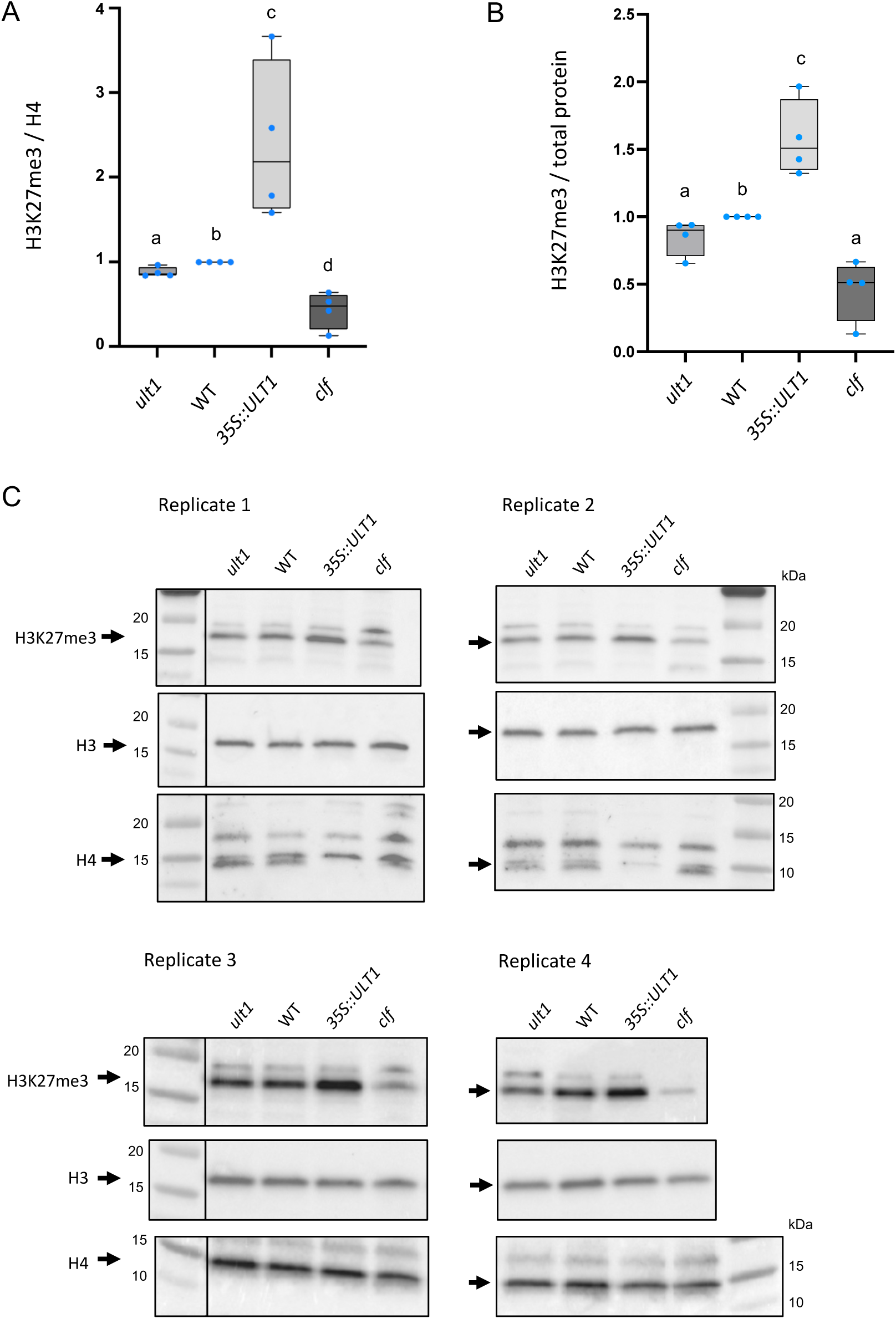
Western blots on 8-day-old seedlings of *ult1-3*, WT, *35S::ULT1 and clf-2*. **(A)** Quantification of H3K27me3 normalized to H4, represented as box plots. Mean was calculated from 4 replicates (which values are indicated by dots) and letters indicate significant differences between samples (Mann-Whitney test, P-value<0.05). **(B)** Quantification of H3K27me3 normalized to total protein in the sample determined using the Qubit Protein Assay Kit (Invitrogen). **(C)** Images of the blot corresponding to the quantifications shown in Fig. 1F and in Suppl. Fig. S4A. Replicate 3 and Replicate 4 were run on the same gel.

**Suppl. Figure S5.**
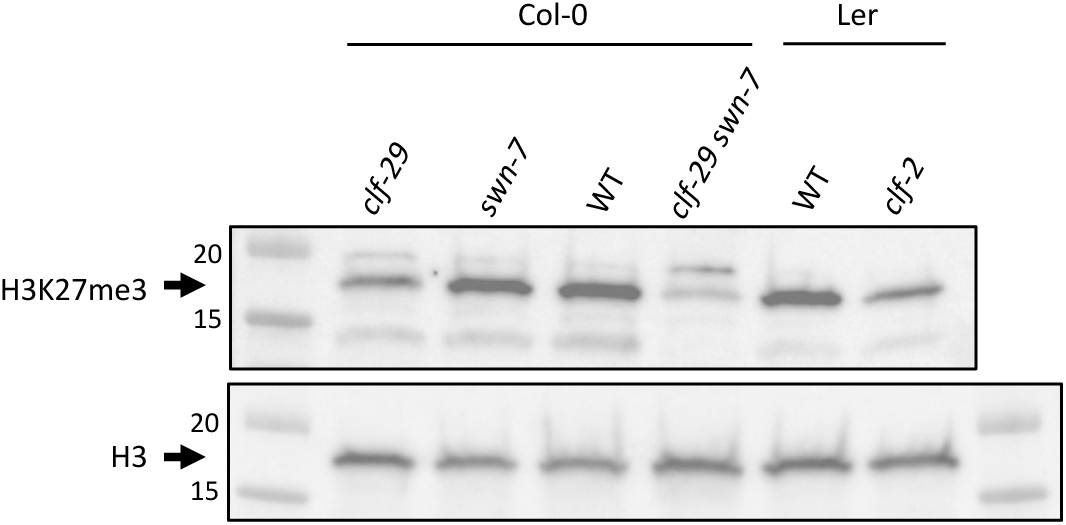
Western blots on 8-day-old seedlings of PRC2 loss-of-function mutants. Assessment of the H3K27me3 antibody used in the study, on nuclear protein extracts from the single mutants *clf-29* and *swn-7* and from the double mutant *clf-29 swn-7* (Col-0 ecotype), along with *clf-2* single mutant (Ler ecotype).

**Suppl. Table S2.** RNA-seq data with the full list of *Arabidopsis thaliana* genes and log2FC between ULT1 lof or ULT1 gof and control WT Ler. Three sheets are displayed for differences between *ult1-3,* or *ult1-2* or *35S::ULT1* and Ler WT. *See excel File*

**Suppl. Table S3.** Lists of genes up and down regulated in the different ULT1lof and ULT1gof lines. Five sheets are displayed in the file, which correspond to, in order: targets targets down-regulated in *ult1-3*, down-regulated in *ult1-2*, down-regulated in *35S::ULT1,* up-regulated in *ult1-3*, up-regulated in *ult1-2*, up-regulated in *35S::ULT1*. P-values were calculated by hypergeometric testing. *See excel File*

**Suppl. Figure S6.**
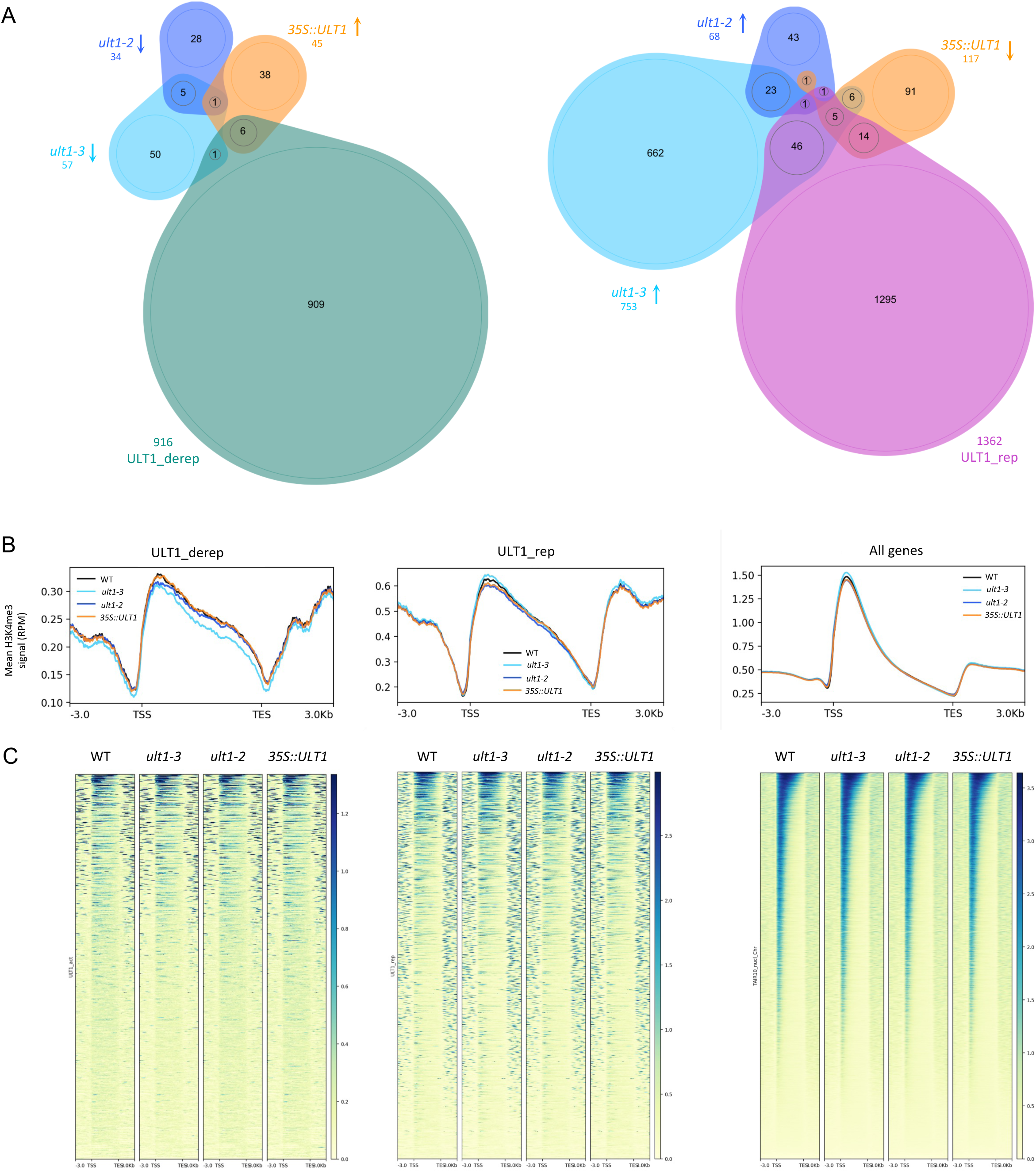
Genome-wide influence of ULT1 on H3K4me3 marks in 8 day-old seedlings (grown in long-day conditions at 18°C day / 16°C night) of ULT1 loss- and gain-of-function lines: WT (Ler wild type), *ult1-2* (EMS line, null mutant), *ult1-3* (T-DNA insertion line, null mutant) and 35S::ULT1 (ectopic surexpressor). **(A)** Venn-diagram showing overlaps between populations of genes (left) with significantly reduced H3K4me3 in *ult1-2* and *ult11-3*, significantly elevated H3K4me3 in 35S::ULT1 and H3K27me3 ULT1_derep genes; and (right) with significantly elevated H3K4me3 in *ult1-2* and *ult1-3*, significantly decreased H3K4me3 in *35S::ULT1* and H3K27me3 ULT1_rep genes. Numbers in the ellipses indicate the number of genes in the respective overlap, numbers under genotypes/gene sets indicate the total number of significantly changed genes found in this genotype/gene set. **(B)** Metagenes for ULT1-derep and ULT1-rep genes: profile of H3K4me3 in WT, *ult1-2*, *ult1-3* and *35S::ULT1* lines. Average distribution of H3K27me3 over ULT1-derep and ULT1-rep genes, and genome-wide. Genes are scaled to the same length (6kb). RPM: reads per million. **(C)** Heatmaps for H3K4me3 distribution for all ULT1-derep and ULT1-rep genes, and genome-wide, ranked from (top line) the highest H3K4me3-marked to(bottom line) the lowest H3K4me3-marked gene in the WT.

**Suppl. Figure S7.**
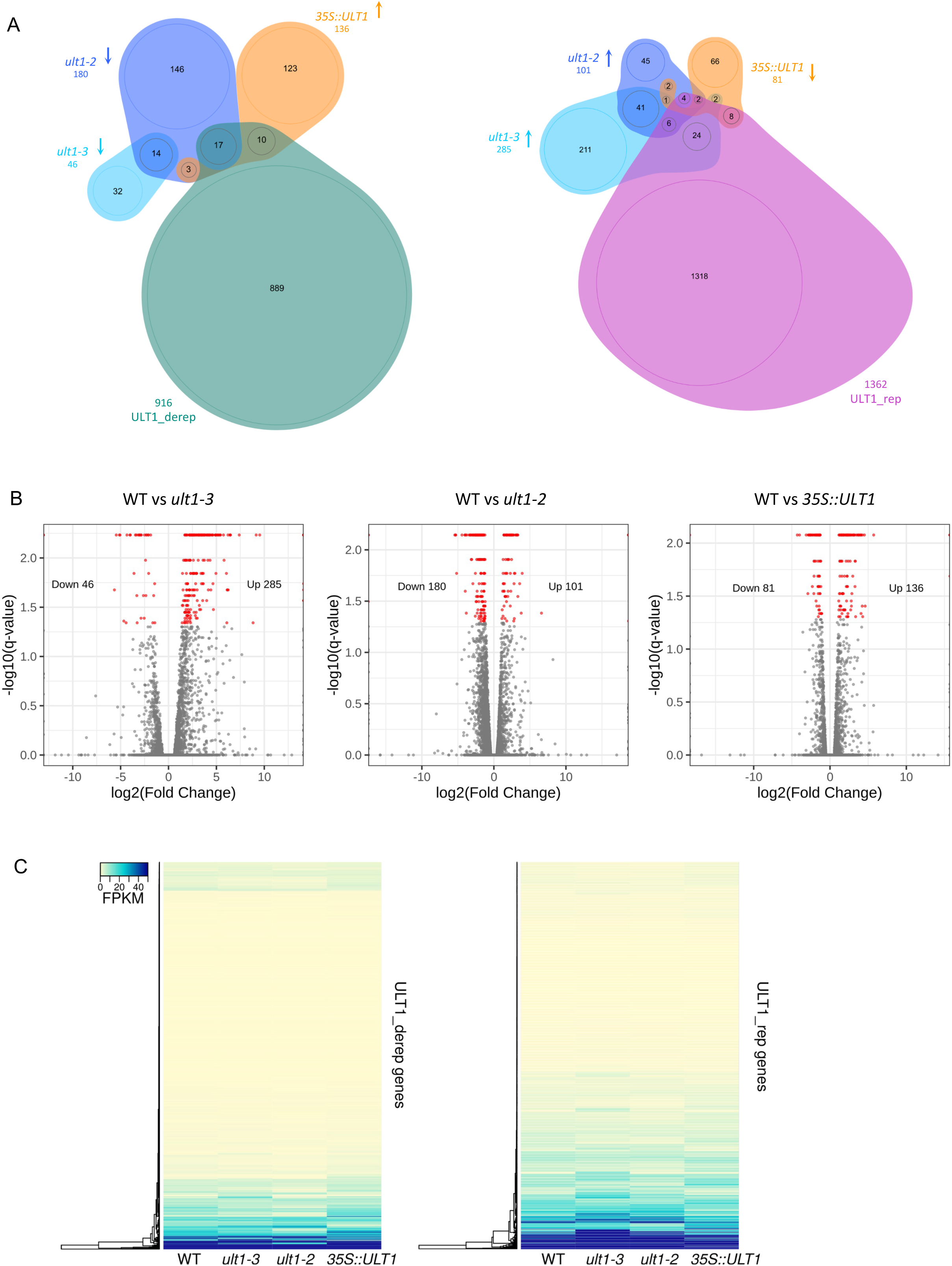
Genome-wide influence of ULT1 on gene expression in 8 day-old seedlings (grown in long-day conditions at 18°C day / 16°C night) **of ULT1 loss- and gain-of-function lines**: WT (Ler wild type), *ult1-2* (EMS line, null mutant), *ult1-3* (T-DNA insertion line, null mutant) and *35S::ULT1* (ectopic surexpressor). **(A)** Venn-diagram showing overlaps between populations of genes (left) with significantly reduced expression in *ult1-2* and *ult11-3*, significantly elevated expression *in 35S::ULT1* and H3K27me3 ULT1_derep genes; and (right) with significantly elevated expression in *ult1-2* and *ult1-3*, signficantly decreased expression in *35S::ULT1* and H3K27me3 ULT1_rep genes. Numbers in the ellipses indicate the number of genes in the respective overlap, numbers under genotypes/gene sets indicate the total number of signi cantly changed genes found in this genotype/gene set. **(B)** Volcano plots showing log2(Fold Change) and –log10(q-value) for all Arabidopsis genes in each genotype comparison. Red dots represent genes showing significant changes in expression between two genotypes, grey dots are genes showing no significant differential expression. **(C)** Heatmaps showing mean expression values (in fragments per kilobase per million reads mapped, FPKM) of three biological replicates in WT, *ult1-2, ult1-3* and *35S::ULT1* genotypes for the H3K27me3 ULT1_derep and H3K27me3 ULT1_rep genes. The maximum expression value was set to 50 FPKM.

**Suppl. Figure S8.**
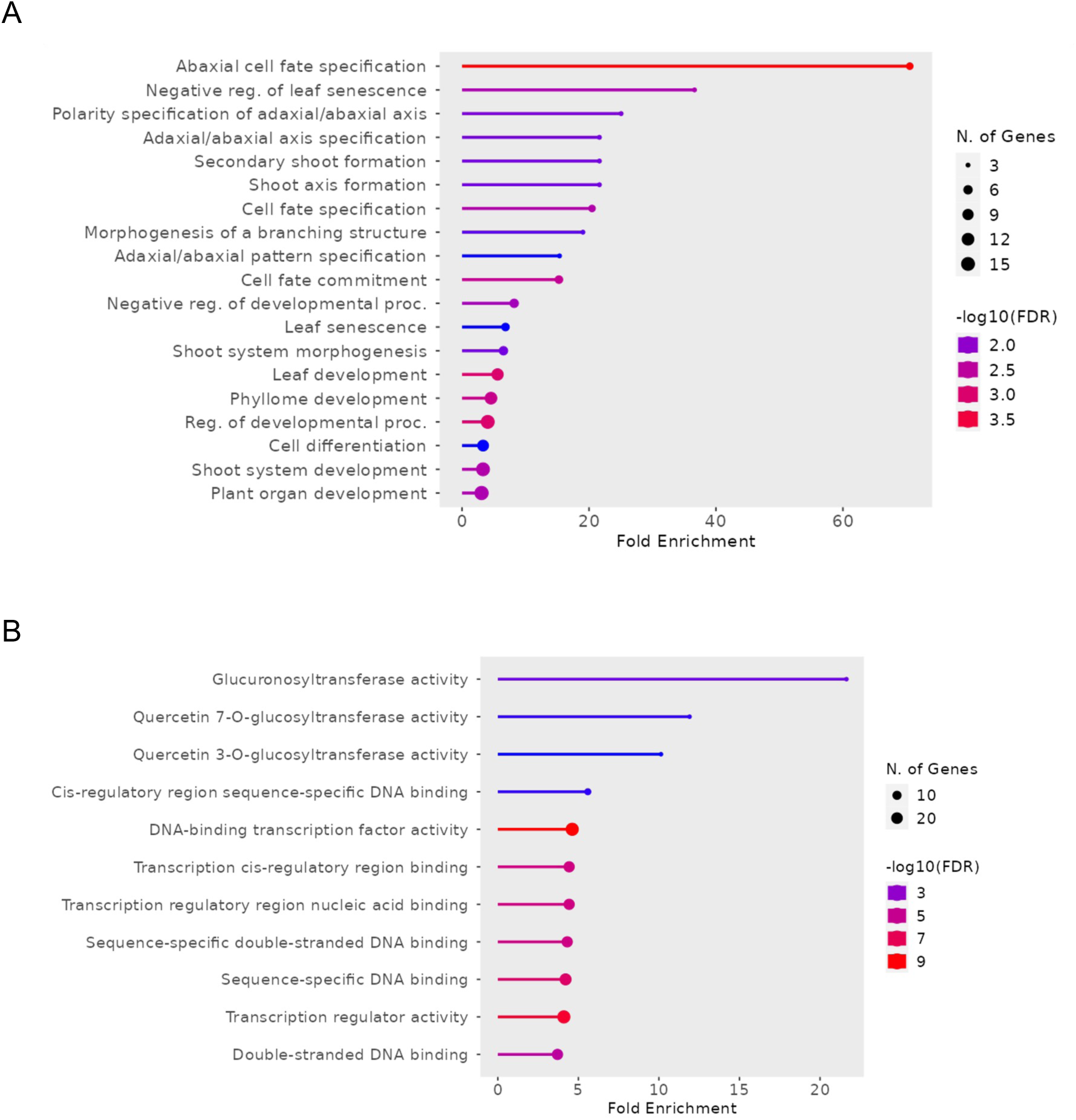
Functional categorisation of genes showing a decrease in H3K4me3 levels in *35S::ULT1*. ShinyGO v0.8 analysis of significantly over-represented Gene Ontology terms among H3K4me3 decreasing genes compared to all H3K4me3 target genes in Ler WT. GO terms are based on **(A)** biological process or **(B)** molecular function.

**Suppl. Table S4.** Overlaps in H3K27me3 changes caused by ULT1 and Histone Methyl Transferases (HMTs) or Histone DeMethylases (HDMTs). Three tables are displayed, that give, for a derepressive function (_derep) or a repressive function (_rep), (i) the Number of target genes in common, (ii) the Percentage of target genes in common (out of n genes targets of <COLUMN>, p % are targets of <LINE>), (iii) the p-value for the above table and (iv) the Sources for the ChIP-seq dataset used. JmJ= the three *Arabidopsis thaliana* H3K27 HDMTs: REF6, ELF6, JMJ13. *See excel File*

**Suppl. Figure S9.**
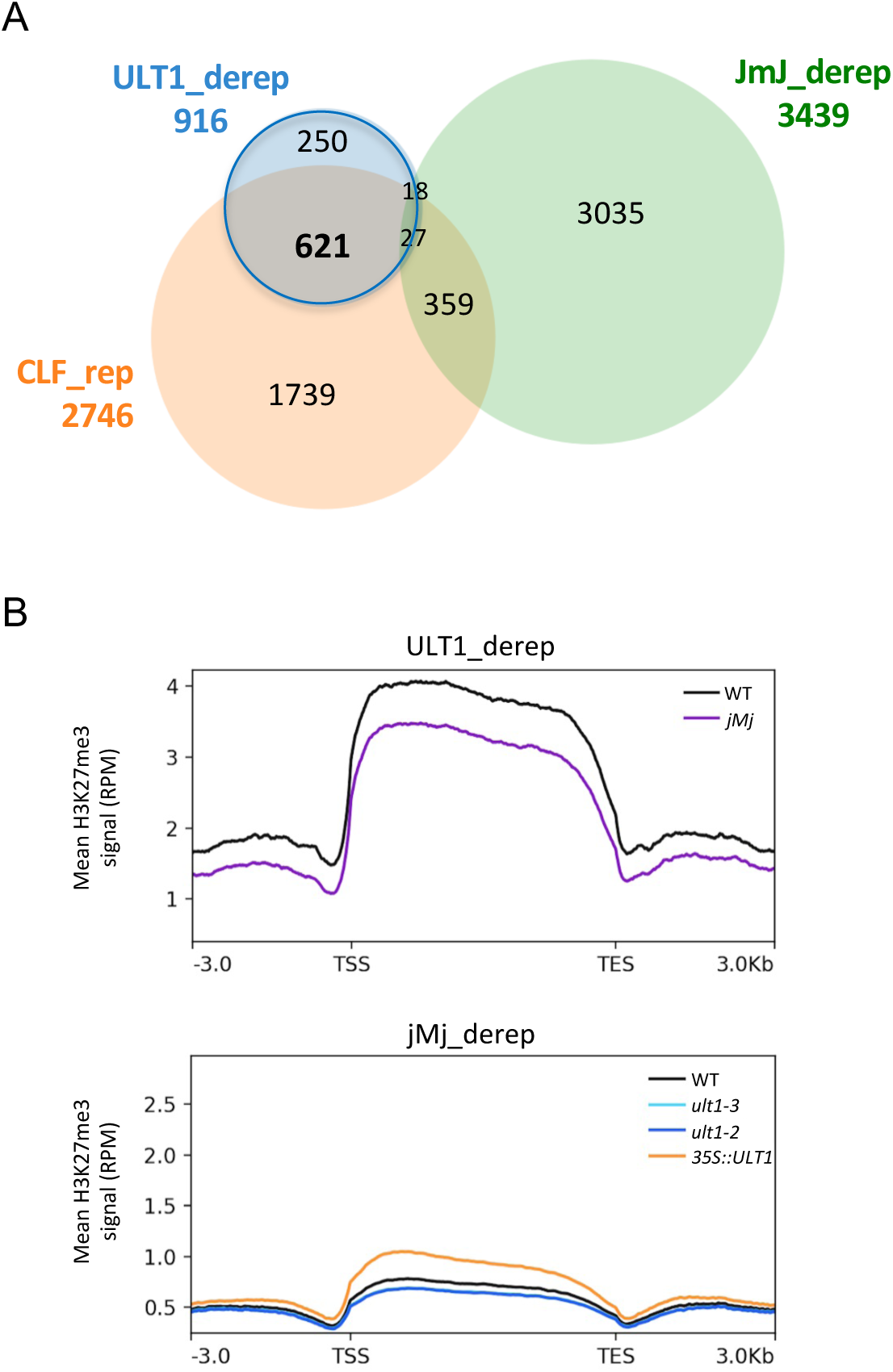
(A) Overlap of genes influenced in their H3K27me3 pattern by ULT1, CLF and the three JmJ H3K27me3 HDMT. Venn-diagram showing overlap between populations of ULT1_derep genes with CLF_rep genes (genes with significantly reduced H3K27me3 in *clf*, raw data from Shu *et al*. 2019) and JmJ derep genes (genes with significantly elevated H3K27me3 in *ref6 elf6 jmj13 = jMj* triple mutant, data from Yan *et al*. 2018). **(B)** Metagenes representing the mean H3K27me3 levels on genes de-repressed by ULT1, as observed in WT and *jMj* mutant (top), and the mean H3K27me3 levels on genes de-repressed by the three jMj, as observed in WT, *ult1-2, ult1-3* and *35S::ULT1* lines (bottom). RPM: reads per million.

**Suppl. Figure S10.**
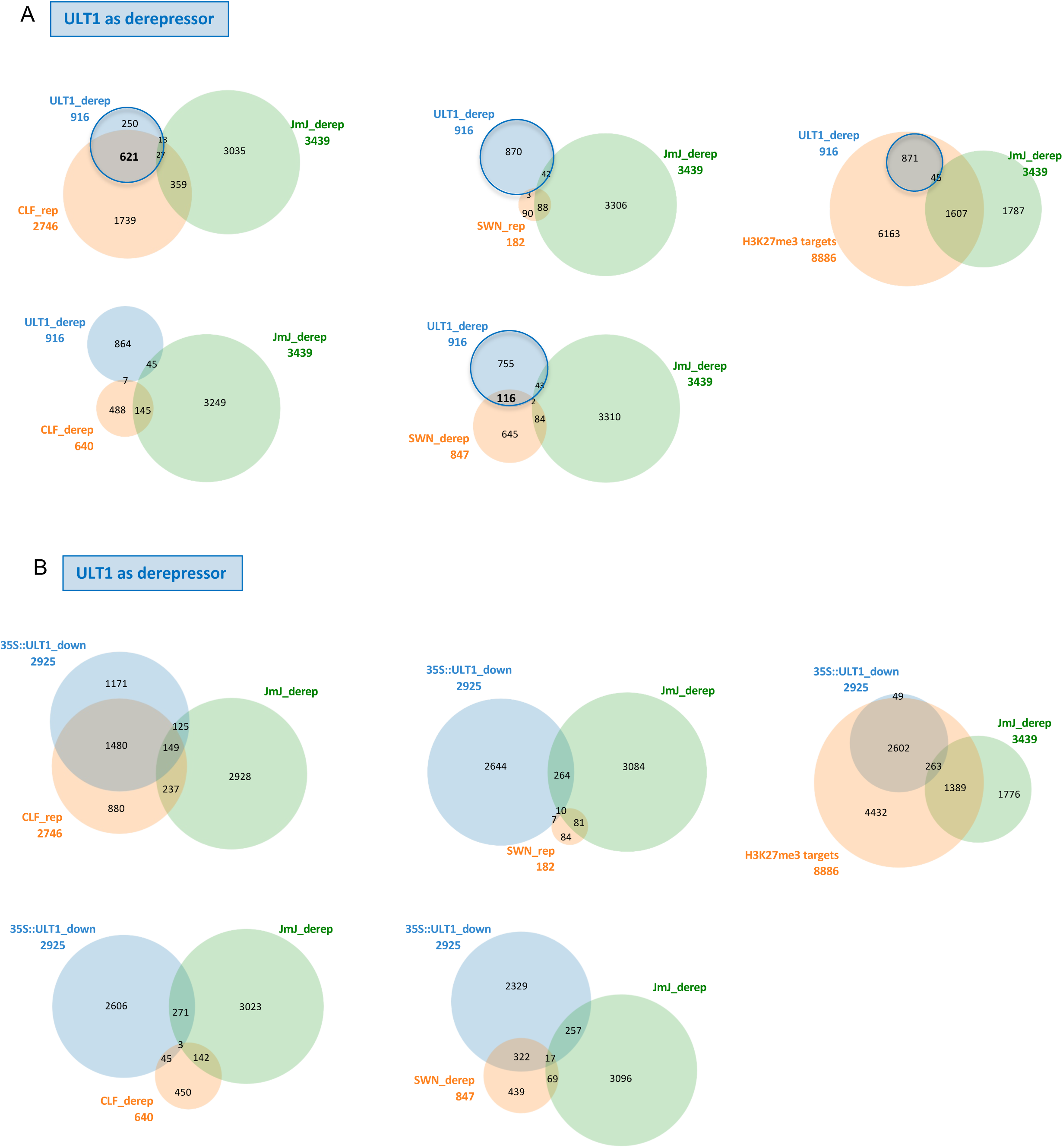
Overlap of genes influenced in their H3K27me3 pattern by ULT1, CLF and the JmJ H3K27me3 HDMT (ULT1_derep). **(A)** Venn-diagram showing overlap between populations of ULT1_derep genes with genes with significantly deregulated H3K27me3 in *clf*, *swn*, or H3K27me3 marked genes (raw data from Shu *et al*. 2019) and JmJ derep genes (genes with significantly elevated H3K27me3 in *ref6 elf6 jmj13* triple mutant, data from Yan *et al*. 2018). **(B)** Venn-diagram showing overlap between populations of 35S::ULT1_down (genes with significantly reduced H3K27me3 in 35S::ULT1) and genes with significantly deregulated H3K27me3 in *clf*, *swn*, or H3K27me3 marked genes and JmJ derep genes (genes with significantly elevated H3K27me3 in *ref6 elf6 jmj13* triple mutant).

**Suppl. Figure S11.**
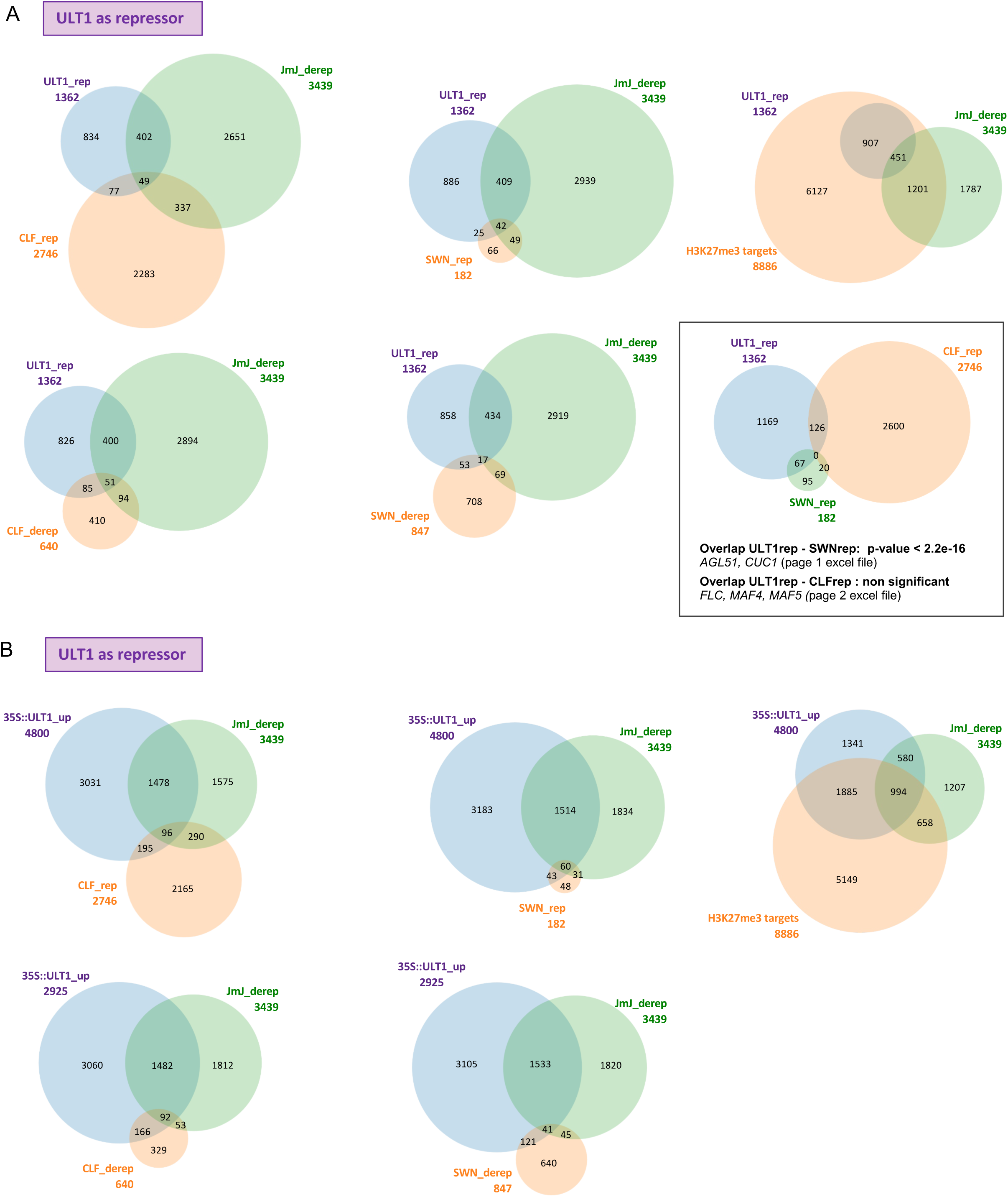
Overlap of genes influenced in their H3K27me3 pattern by ULT1, CLF and the three JmJ H3K27me3 HDMT (ULT1_rep). **(A)** Venn-diagram showing overlap between populations of ULT1_rep genes with genes with significantly deregulated H3K27me3 in *clf*, *swn*, or H3K27me3 marked genes (raw data employed from Shu *et al*. 2019) and JmJ derep genes (genes with significantly elevated H3K27me3 in *ref6 elf6 jmj13* triple mutant, data from Yan *et al*. 2018). **(B)** Venn-diagram showing overlap between populations of 35S::ULT1_up (genes with significantly increased H3K27me3 in *35S::ULT1*) and genes with significantly deregulated H3K27me3 in *clf*, *swn*, or H3K27me3 marked genes and JmJ derep genes (genes with significantly elevated H3K27me3 in *ref6 elf6 jmj13* triple mutant).

**Suppl Figure S12.**
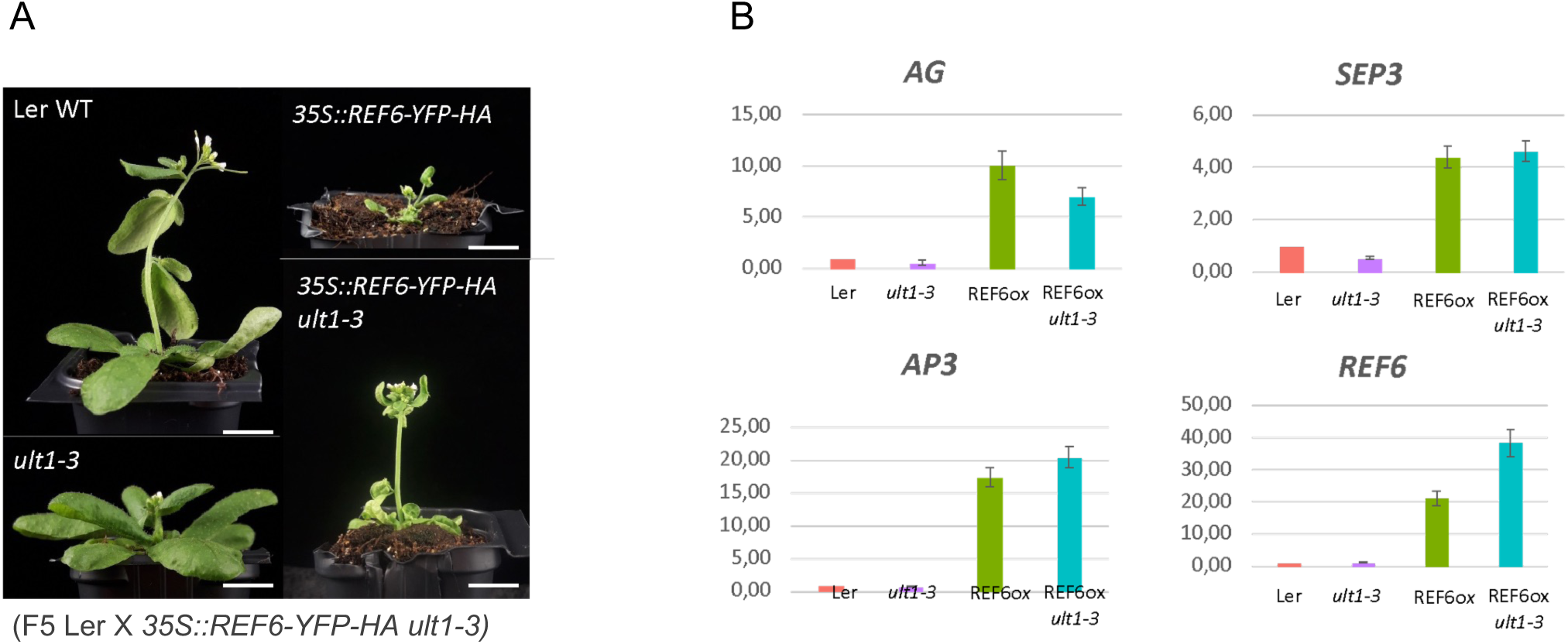
REF6 is dispensable for ectopic ULT1-induced phenotypes and de-repression of MADS floral genes. **(A)** Pictures of 28 day-old Ler plants (WT, *35S::REF6-YFP-HA*, *ult1-3* mutant, *35S::REF6-YFP-HA ult1-3),* indicate that *35S::REF6-YFP-HA* ectopic expression phenotypes do not depend on ULT1 function. **(B)** Relative expression levels of the homeotic floral genes *AG, AP3* and *SEP3,* along with *REF6*, in 8-day-old seedlings, assessed by RT-qPCR. Expression levels of target genes were normalized with the reference gene *MON1*. Error bars correspond to standard errors for 3 technical replicates. Similar results were obtained for 2 biological replicates.

**Suppl. Table S5.** List of proteins copurifying with ULT1 in AP-MS on Arabidopsis cell cultures. The list corresponds to proteins detected by LC-MS/MS in a GS-ULT1 pulldown (found at least in two out of three replicates, Mascot Score >80). The protein list of two control pulldowns of unfused GS (“GS-only”) was used to filter out background contaminants. In addition, a published list (van Leene et al. 2015) of common AP-MS contaminants was used for filtering. *See excel File*

**Suppl. Figure S13.**
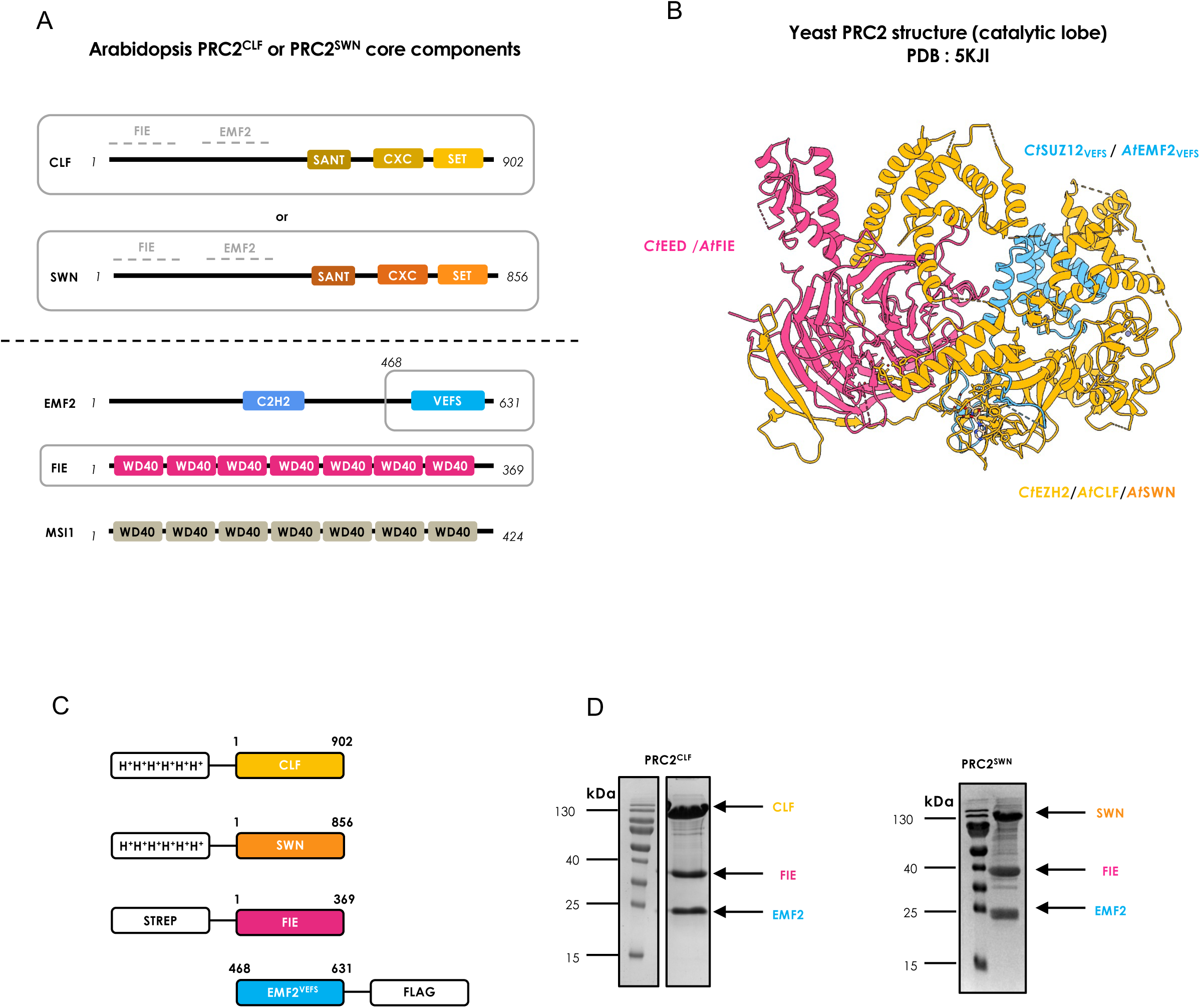
Expression strategy for the catalytic lobes of the Arabidopsis PRC2^CLF^ and PRC2^SWN^ complexes. **(A)** Schematic representation of the PRC2 complex subunits in *Arabidopsis thaliana*. The catalytic lobe of the PRC2 complex consists of a methyltransferase which is either CLF (yellow) or SWN (orange), the WD40 domain protein FIE (pink) and the VEFS domain of EMF2 (blue). The subunits forming the ternary complex are represented by a grey frame. Areas of interaction of FIE and EMF2 with CLF or SWN are represented by gray bars. The domains of each protein identified using Prosite analysis are represented by colored rectangles. **(B)** Structure of the catalytic lobe of PRC2 from *Chaetomium thermophilum* determined by X-ray crystallography; PDB: 5KJI (Jiao and Liu, Science 2015) that was used to design the constructs of each subunit. The subunits are represented using the same color code as in (A). **(C)** Schematic representation of the constructs made for each subunit of the PRC2 complex. The coding sequences for CLF_1-902_ (or SWN_1-856_), FIE_1-369_ and EMF2_468-631_ were cloned into expression vectors for production of recombinant proteins in fusion with His_6_, Strep or Flag tags, respectively. The complex purification tags associated with each subunit and their positions (N-terminal or C-terminal) are indicated. **(D)** SDS-PAGE of the PRC2^CLF^ and PRC2^SWN^ complexes after purification by size exclusion chromatography.

**Suppl. Figure S14.**
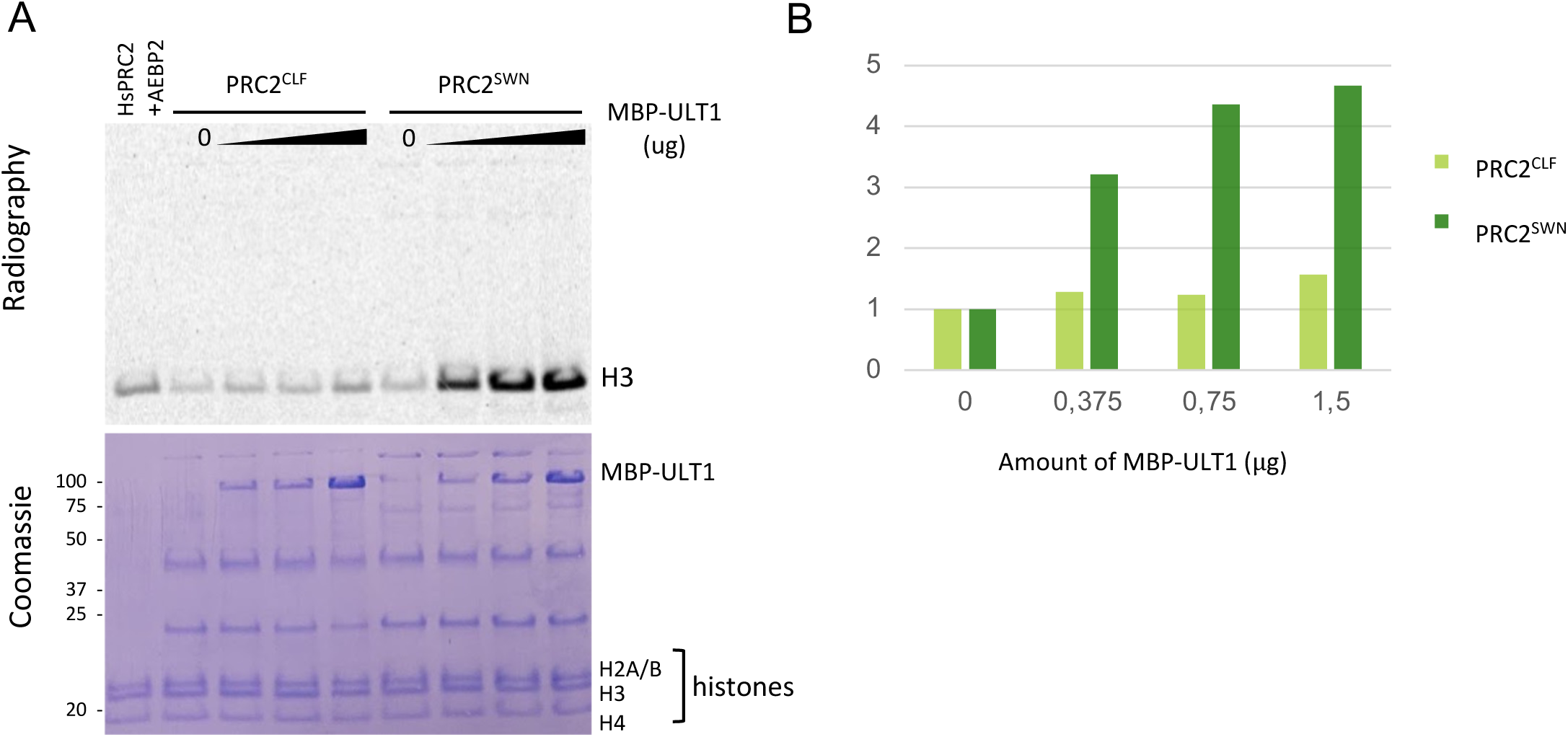
*In vitro* HMT activity tests show ULT1-mediated increase of the PRC2^SWN^ activity but not of the PRC2^CLF^ activity on recombinant nucleosomes. **(A)** Top panel shows autoradiography of a SDS-PAGE gel after incubation of PRC2^CLF^ or PRC2^SWN^ (1.5ug of complex) without or with ULT1 in increasing amounts (0, 0.375, 0.75, 1.5ug). The HsPRC2 complex (0.5ug) was loaded in the first lane after incubation with the cofactor AEBP2, for positive control. Bottom panel shows Coomassie staining of the same SDS-PAGE. **(B)** Graph showing relative quantifications of HMT activities for PRC2^CLF^ and PRC2^SWN^ incubated with MBP-ULT1.

**Suppl. Figure S15.**
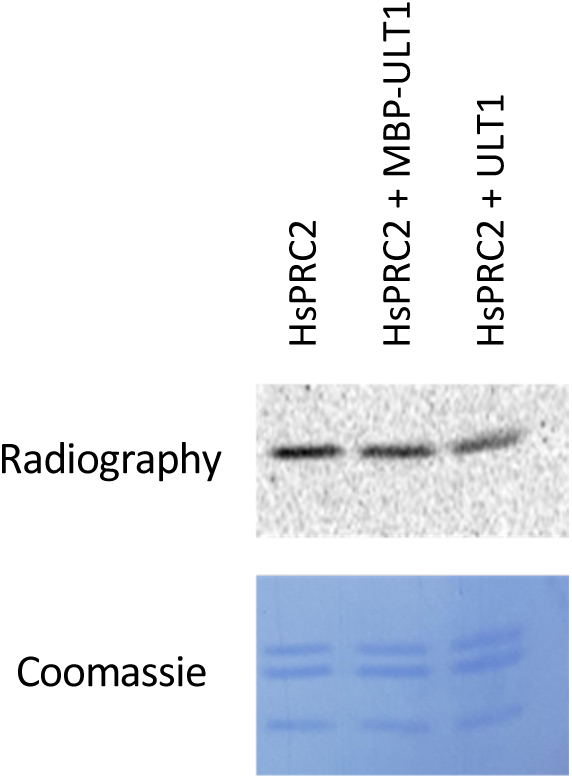
*In vitro*, ULT1 does not impact the activity of the HsPRC2 EZH2 complex on recombinant nucleosomes.

**Suppl. Figure S16.**
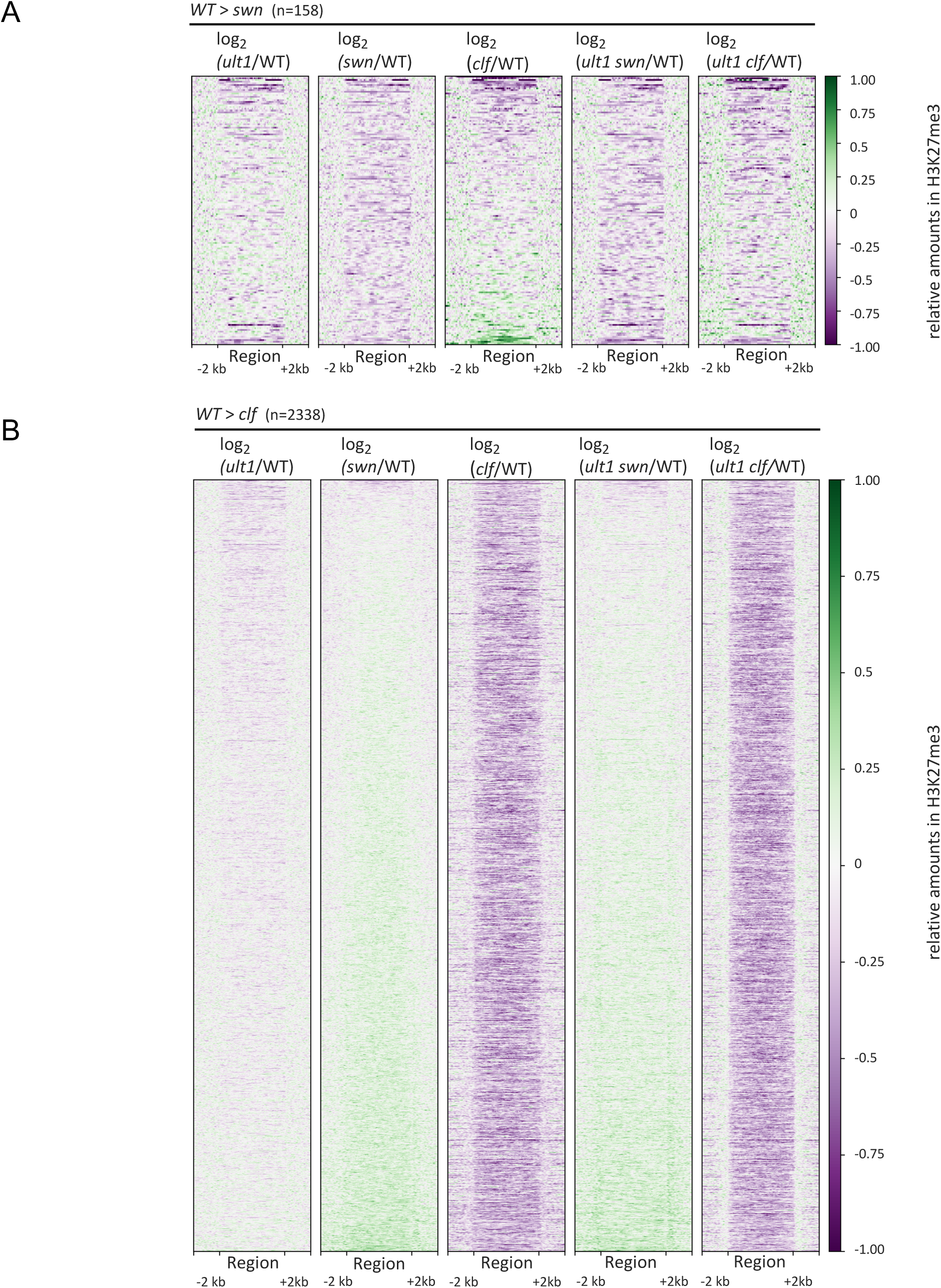
Heatmaps showing differences in H3K27me3 abundance in *ult1*, *swn*, *clf*, *ult1swn* and *ult1clf* compared to WT. Areas displayed contain (A) regions with reduced H3K27me3 in *swn* compared to WT (n= 158 regions, p<0.05) sorted by their difference in *clf*, and (B) regions with reduced H3K27me3 in *clf* compared to WT (n= 2338 regions, p<0.0001) sorted by their difference in *swn*. Note that for determination of regions with reduced H3K27me3 in *swn* a more lenient p-value was used to provide a more general overview of regions with reduced H3K27me3 and not limit this analysis to the only 67 regions found at the more strict p-value of 0.0001. Differences are depicted as log2 fold-change values with respect to WT in heatmap by a color gradient ranging from green for positive values and higher H3K27me3 abundance in the WT to purple for negative values indicating higher mark abundance in the respective mutant. Each line represents one region, all regions are scaled to the same length. At the 158 regions with reduced H3K27me3 in the *swn* single mutant (A), a similar pattern is observed in *swn* and *ult1 swn* for H3K27me3 changes compared to WT. Interestingly, this set contains a few regions showing a strong enhancement of H3K27me3 abundance in the *clf* mutant, likely caused by an overly active PCR^SWN^ complex at these regions. Consistent with ULT1 assisting SWN in its HMT activity, this enhancement of H3K27me3 signal is largely lost in the *ult1clf* double mutant. This observation is not mirrored at the 2338 regions with reduced H3K27me3 abundance in the *clf* single mutant (B): while mainly conserved in the *ult1 clf* double mutant, a large fraction of regions show higher H3K27me3 signal in the *swn* single mutant, and this enhanced signal is not reduced in the *ult1 swn* double mutant. This observation hints at a stronger activity of CLF at these loci when SWN is absent and this strong CLF activity is still present when ULT1 is absent.

**Suppl. Figure S17.**
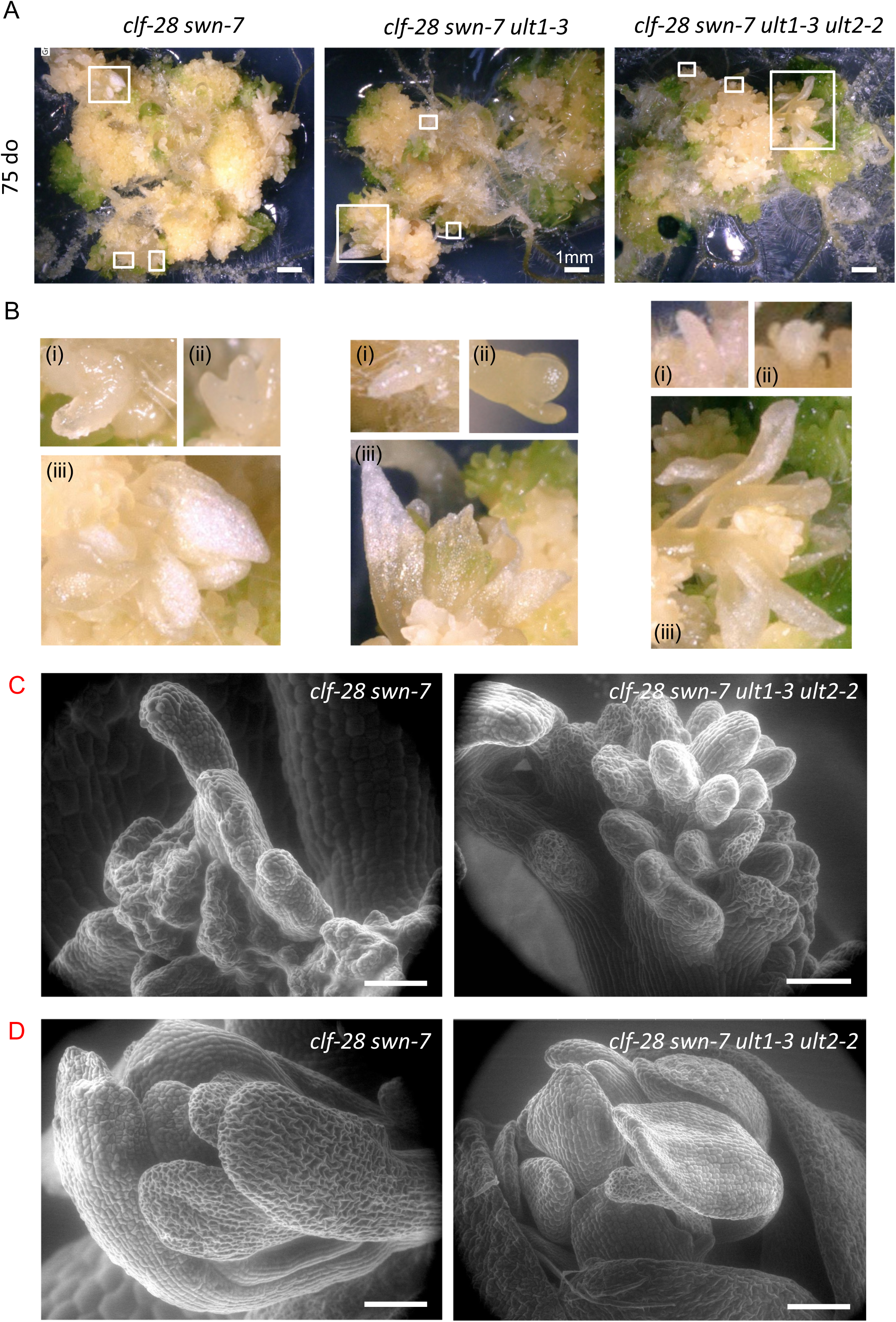
*ult* loss-of-function does not complement sporophytic *prc2* mutant phenotypes. **(A)** Pictures of double *clf-28 swn-7, triple clf-28 swn-7 ult1-3* and quadruple *clf-28 swn-7 ult1-3 ult2-2* mutants grown on MS agar plates for 2.5 months. All genotypes grow as callus-like tissue with random organs arising in a disorganised phyllotaxy. **(B)** Magnified views of insets drawn in panels shown in (A), showing ectopic (i) roots and hair roots, (ii) somatic embryos or shoot apex-like structure, and (iii) developed shoots with some homeotic conversions to floral organ types. Scale bar: 1mm. **(C-D)** SEM images of *clf-28 swn-7* and *clf-28 swn-7 ult1-3 ult2-2* loss-of-function lines show stigmatic papillae-type outgrowths and growth of flower-like structures. Pictures of double *clf-28 swn-7* and quadruple *clf-28 swn-7 ult1-3 ult2-2* mutants grown on MS agar plates for 3 weeks. **(C)** Ectopic stigmatic papillae-like outgrowths. **(D)** Organs arranged in flower-like structures. Scale bar: 100um.

**Suppl. Figure S18.**
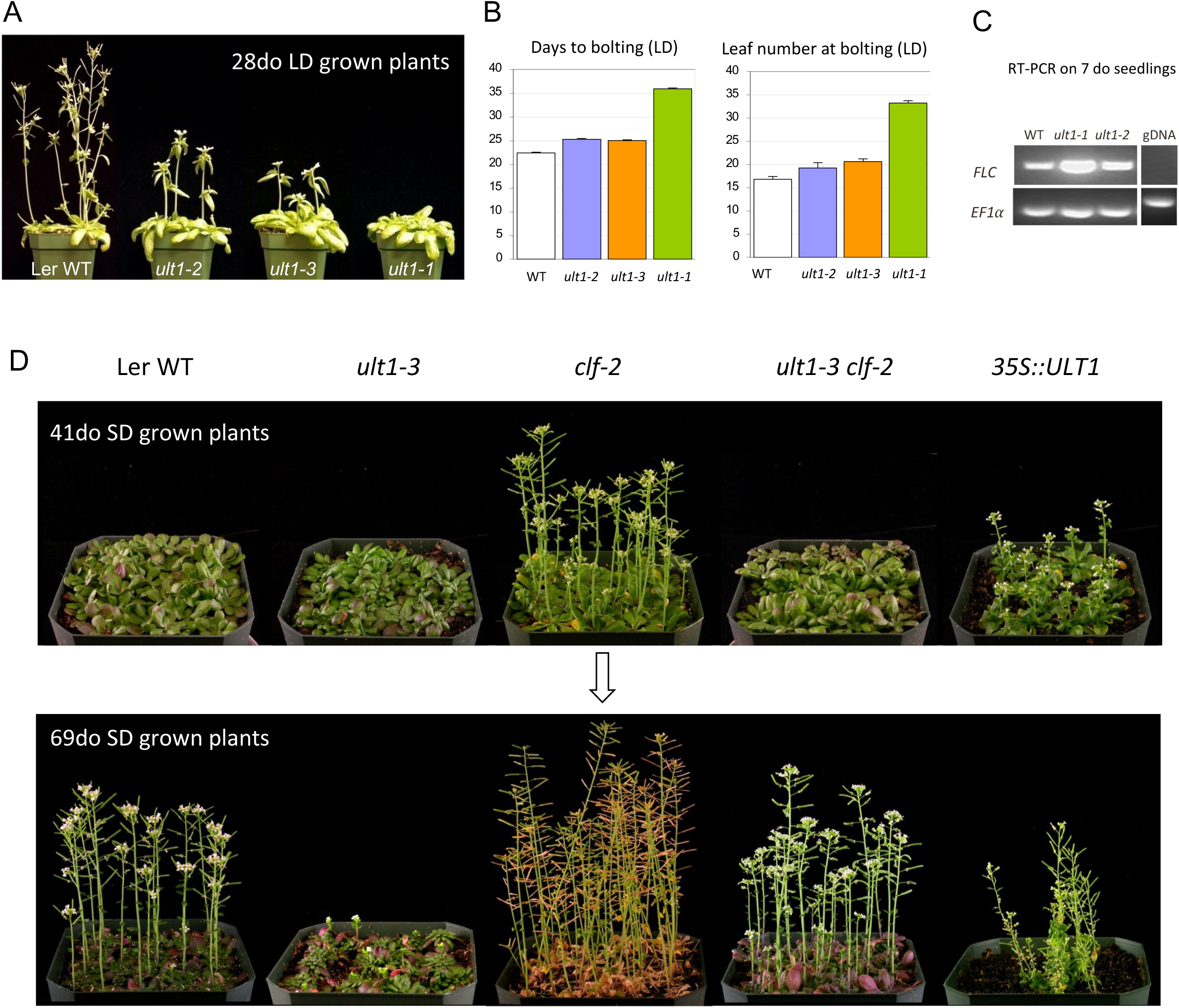
Flowering time of ULT1 lof and gof plants in the Ler ecotype, in LD and SD. **(A-C)** Plants grown in LD condition. (A) Representative 28 day-old Ler WT, *ult1-2, ult1-3* and *ult1-1* mutant plants grown at 20 °C in long days. **(B)** Number of days to bolting and number of rosette leaves were scored as proxy for the flowering time. The previousley reported semi-dominant *ult1-1* mutant allele exibits a more dramatic late flowering phenotype than the null *ult1-2* and *ult1-3* alleles. **(C)** RT-PCR performed on 7 day-old plants reveals a high upregulation of *FLC* expression. **(D)** Plants grown in SD condition. Representative Ler WT, *ult1-3, clf-2, ult1-3 clf-2* and *35S::ULT1* plants grown at 20 °C in short days were pictured at 41 and 69 days old.

**Suppl. Figure S19.**
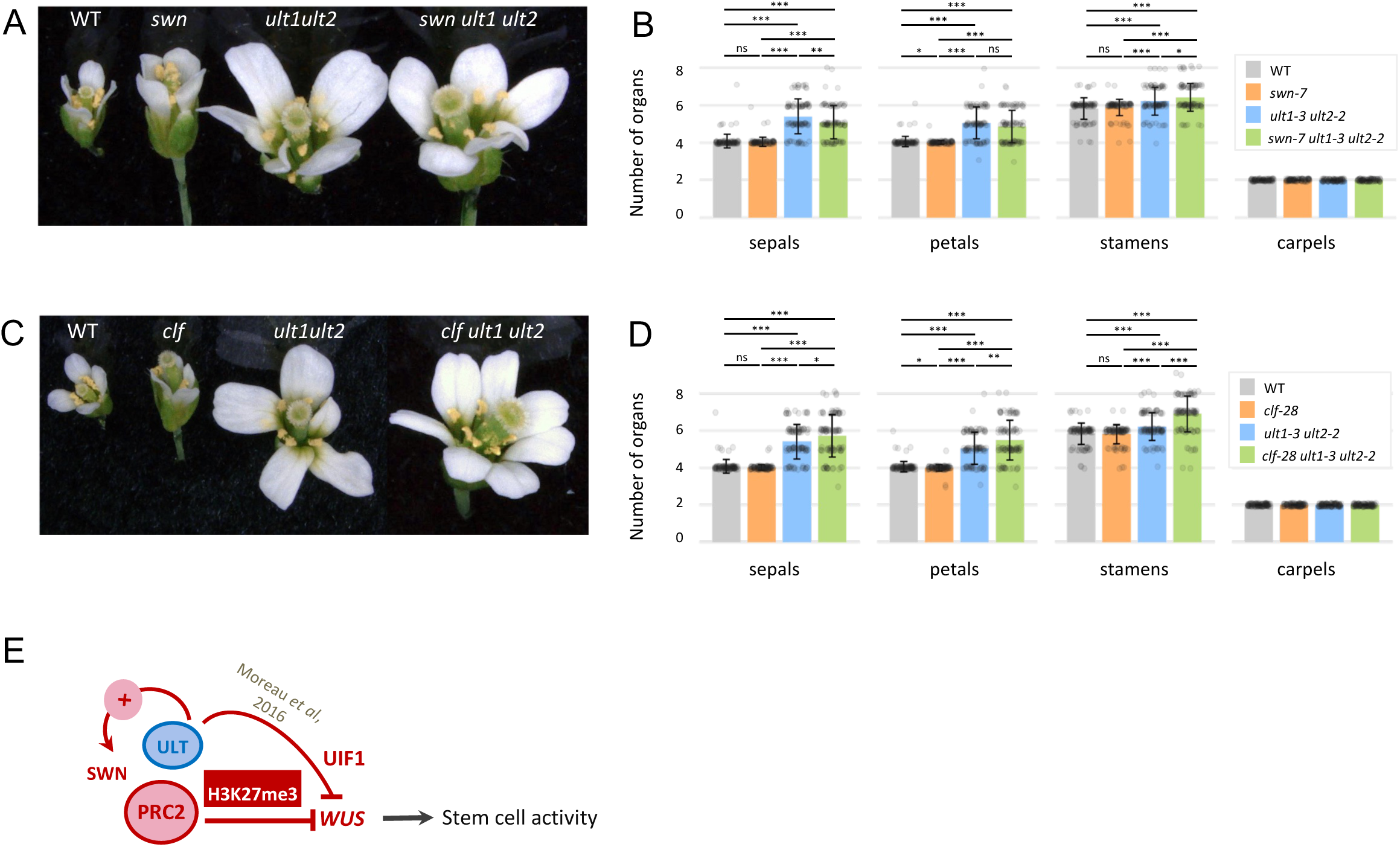
Genetic interactions between ULT and sporophytic PRC2 functions for flower meristem maintenance. **(A)** Representative flowers of Col-0 WT, *swn-7, ult1-3 ult2-2* and *swn-7 ult1-3 ult2-2* plants. **(B)** Organ numbers in each floral whorl, for all above mentioned genotypes. Flowers of *ult1 ult2 swn* triple mutant produce the same number of organs as flowers of *ult1 ult2* double mutants, for each whorl. **(C)** Representative flowers of Col-0, *clf-28, ult1-3 ult2-2* and *clf-28 ult1-3 ult2-2* mutant plants. **(D)** Organ numbers in each floral whorl, for all above mentioned genotypes. Flowers of *ult1 ult2 clf* triple mutant produce more floral organs than *ult1 ult2* double mutants, indicating a synergy between ULT and CLF pathways. In B,C: n (number of counted flowers) = 100 for all genotypes except for *swn-7 ult1-3 ult2-2* (n=99). p-values are as follows: * *p* < 0.05, ** *p* < 0.01, *** *p* < 0.001, ns = not significant. **(E)** Working model for stem cell activity at the floral meristem, and for *WUS* regulation by ULT1 and PRC2. ULT1 functions together with SWN to promote *WUS* repression. Moreover, another pathway could be considered for ULT, independent of PRC2, involving a direct interaction with the UIF1 transcription factor which has a repressive function on *WUS* expression (Moreau *et al*., 2016).

**Suppl. Table S6.** List and sequences of primers used in this study. *See excel File*

